# A systematic analysis of metabolic pathways in the human gut microbiota

**DOI:** 10.1101/2021.02.25.432841

**Authors:** Victòria Pascal Andreu, Hannah E. Augustijn, Lianmin Chen, Alexandra Zhernakova, Jingyuan Fu, Michael A. Fischbach, Dylan Dodd, Marnix H. Medema

## Abstract

The gut microbiota produce hundreds of small molecules, many of which modulate host physiology. Although efforts have been made to identify biosynthetic genes for secondary metabolites, the chemical output of the gut microbiome consists predominantly of primary metabolites. Here, we systematically profile primary metabolic genes from the gut microbiome, identifying 19,885 gene clusters in 4,240 high-quality microbial genomes. We find marked differences in pathway distribution among phyla, reflecting distinct strategies for energy capture. These data explain taxonomic differences in short-chain fatty acid production and suggest a characteristic metabolic niche for each taxon. Analysis of 1,135 subjects from a Dutch population-based cohort shows that the level of 14 microbiome-derived metabolites in plasma is almost completely uncorrelated with the metagenomic abundance of the corresponding biosynthetic genes, revealing a crucial role for pathway-specific gene regulation and metabolite flux. This work is a starting point for understanding differences in how bacterial taxa contribute to the chemistry of the microbiome.

---

The pathways encoding the production of microbial metabolites are often physically clustered in the genome, in regions known as metabolic gene clusters (MGCs). Current tools for computational prediction of metabolic pathways focus on gene clusters for natural product biosynthesis (*1*) or generic primary metabolism (*2, 3*). Here, we introduce a new algorithm, gutSMASH, to profile known and predicted novel primary metabolic gene clusters from the gut microbiome. We use this tool to perform a systematic analysis of primary metabolic gene clusters in bacterial strains from the gut microbiome, and identify the prevalence and abundance of each of these pathways across a large population-based cohort.

Algorithms that identify physically clustered genes have become a mainstay of bacterial pathway identification; taking into account the conserved physical clustering of genes prevents false positive hits based on sequence similarity alone. This principle has been widely applied in the field of natural product biosynthesis, e.g. in antiSMASH (*1*), which predicts biosynthetic gene clusters (BGCs) by detecting physically clustered protein domains using profile hidden Markov Models (pHMMs). Here, *we* tailored this gene cluster detection framework to detect MGCs involved in primary metabolism and bioenergetics.

As a starting point, we constructed a dataset of 51 primary metabolic pathways from the gut microbiome with biochemical or genetic literature support (including MGCs as well as pathways encoded by a single gene) and identified core enzymes (i.e., required for pathway function) to serve as a signature for the detection rules (Figure 1, Table S1; see *Methods* for details). To more accurately predict MGCs of interest, we performed three computational procedures. First, for core enzymes belonging to 12 of the protein superfamilies that are known to catalyze diverse types of reactions and were most commonly found across a wide range of pathways, we constructed phylogenies and used them to create clade-specific pHMMs to detect specific subfamilies (see SI results *Phylogenetic analysis of protein superfamilies to identify pathway-specific clades*). Second, we designed pathway-specific rules for each MGC type in our dataset (see *Methods*). These rules were validated and optimized by detailed visual inspection and analysis of MGC sequence similarity networks made using BiG-SCAPE (*4*), generated from gutSMASH results on a set of 1,621 microbial genomes (Online Data: https://gutsmash.bioinformatics.nl/help.html#Validation); see SI results *Validation of gutSMASH detection rules by evaluating their predictive performance*) (Table S2&S3). Third, despite the fact that most specialized primary metabolic pathways are encoded in MGCs, there are also single-protein pathways that are in charge of the secretion of key specialized primary metabolites in the gut microbial ecosystem, such as serine dehydratase, which produces ammonia and pyruvate from serine (*5*). For this reason, we also built 10 clade-specific pHMMs to detect these (see Methods section *Assessing single-protein pathway abundance within representative human gut bacteria*). The above procedures led to the design of a robust set of detection rules to identify both known and putative MGCs that are potentially relevant for metabolite-mediated microbiome-associated phenotypes.

**Figure 1:**
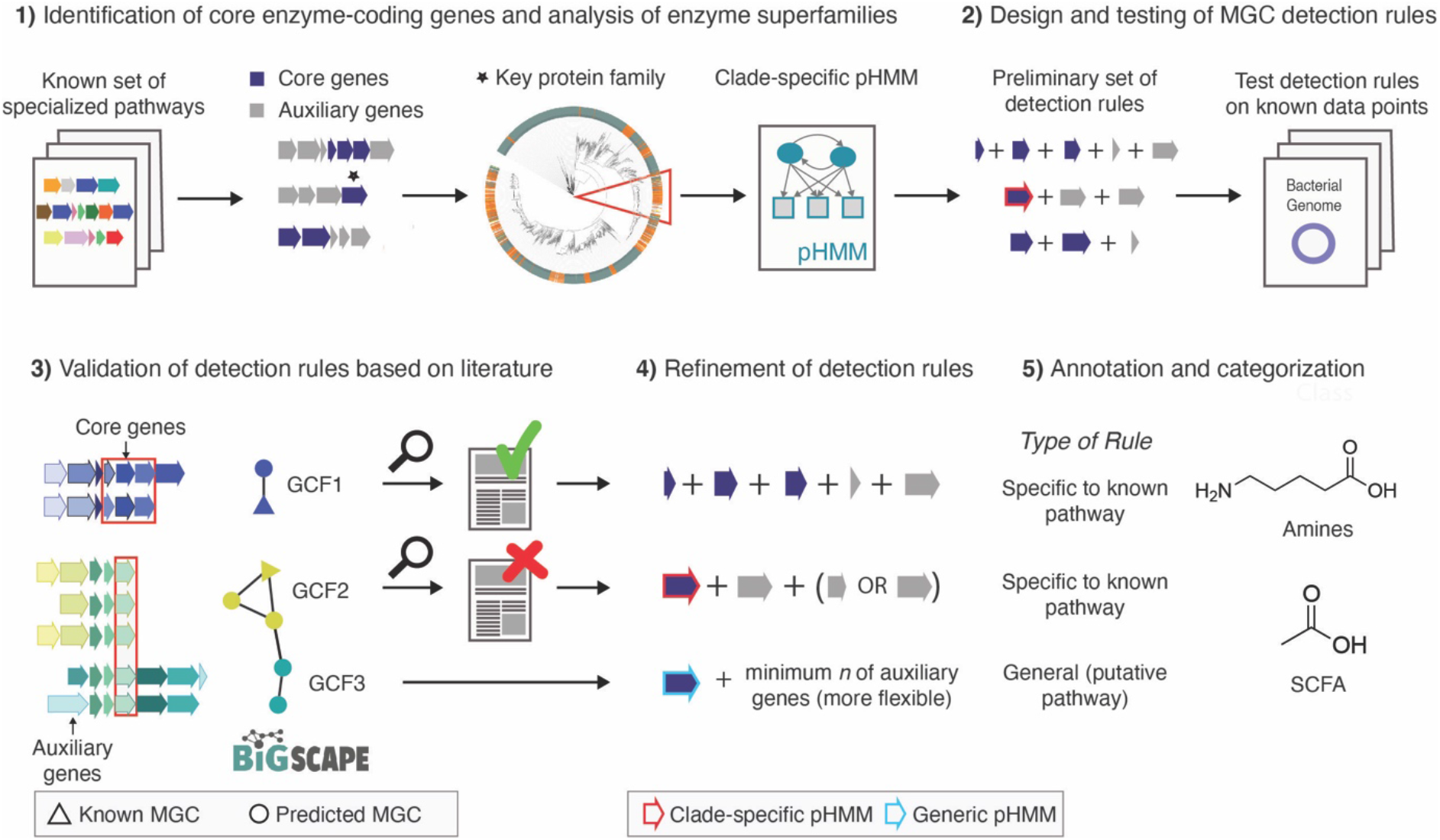
Development and design of detection rules for gutSMASH. (1) A set of known and characterized MGC-encoded pathways were curated from the literature. Protein domains were identified across all MGCs and core enzymatic domains were manually identified. For enzymatic domains belonging to broad multifunctional enzyme families, protein superfamily phylogenies were built to create clade-specific pHMMs. (2) These domains were incorporated in the initial detection rules. The detection rules were run on a test set, and all the MGC predicted by the same rule were grouped together and (3) run through BiG-SCAPE, which grouped the MGCs into gene cluster families (GCFs). (4) Based on literature analysis of GCF members, detection rules were manually fine-tuned to either include or exclude MGC architectures that were either related to specialized primary metabolism or not. (5) Finally, fine-tuned detection rules were annotated and categorized into different MGC classes based on their metabolic end products.

To profile the metabolic capacity of strains from the human gut microbiome, we selected a set of 4,240 unique high-quality reference genomes consisting of 1,520 genomes from the Culturable Genome Reference (CGR) collection (*6*), 2,308 genomes from the Microbial Reference Genomes collection of the Human Microbiome Project (HMP) consortium (*7*) and 414 additional genomes from the class Clostridia to account for their metabolic versatility (*8*) (Table S4). We refrained from including metagenome-assembled genomes in this analysis, as they often lack the taxon-specific genomic islands (*9*) on which many specialistic metabolic functions are encoded. In total, gutSMASH predicted 19,885 MGCs across these genomes that are clear homologues of MGCs for our set of known pathway types (See Methods: *Evaluating the functional potential of the human microbiome using gutSMASH*).

The combined results of the gutSMASH MGC scanning and the single-protein pHMM detection across the three reference collections provide unique insights into the metabolic traits encoded by the genomes of human gut bacteria. While some genera harbor a small set of highly conserved pathways, (e.g., *Akkermansia, Faecalibacterium*), other genera contain much larger interspecies differences (Figure 2A). The genus *Clostridium* displays remarkable metabolic versatility, with 42 distinct metabolic pathways present across members of this genus (Figure 2A). Clostridial strains that are indistinguishable by 16S sequencing often harbor distinct gene cluster ensembles (Suppl. Figure 1), suggesting that specialization in primary metabolism leads to functional differentiation even among closely related strains.*Clostridium* is a clear outlier: by comparison, the next most numerous set of metabolic pathways are found within the Enterobacteriaceae (e.g., *Salmonella, Escherichia, Enterobacter, and Klebsiella*) with 22-25 metabolic pathways. Intriguingly, many of the metabolic pathways encoded by *Clostridium* and members of the Enterobacteriaceae are non-overlapping (with 23/42 *Clostridium* pathways not being identified among Enterobacteriaceae), highlighting the distinct metabolic strategies these microbes employ within the gut (Figure 2A). The *Bacteroides*,Actinobacteria (*Eggerthella* and *Collinsella*) and Verrucomicrobia (*Akkermansia*) harbor a more restricted set of primary metabolic pathways, likely reflecting versatility in upstream components of their metabolism (i.e., glycan foraging and other forms of substrate utilization).

**Figure 2:**
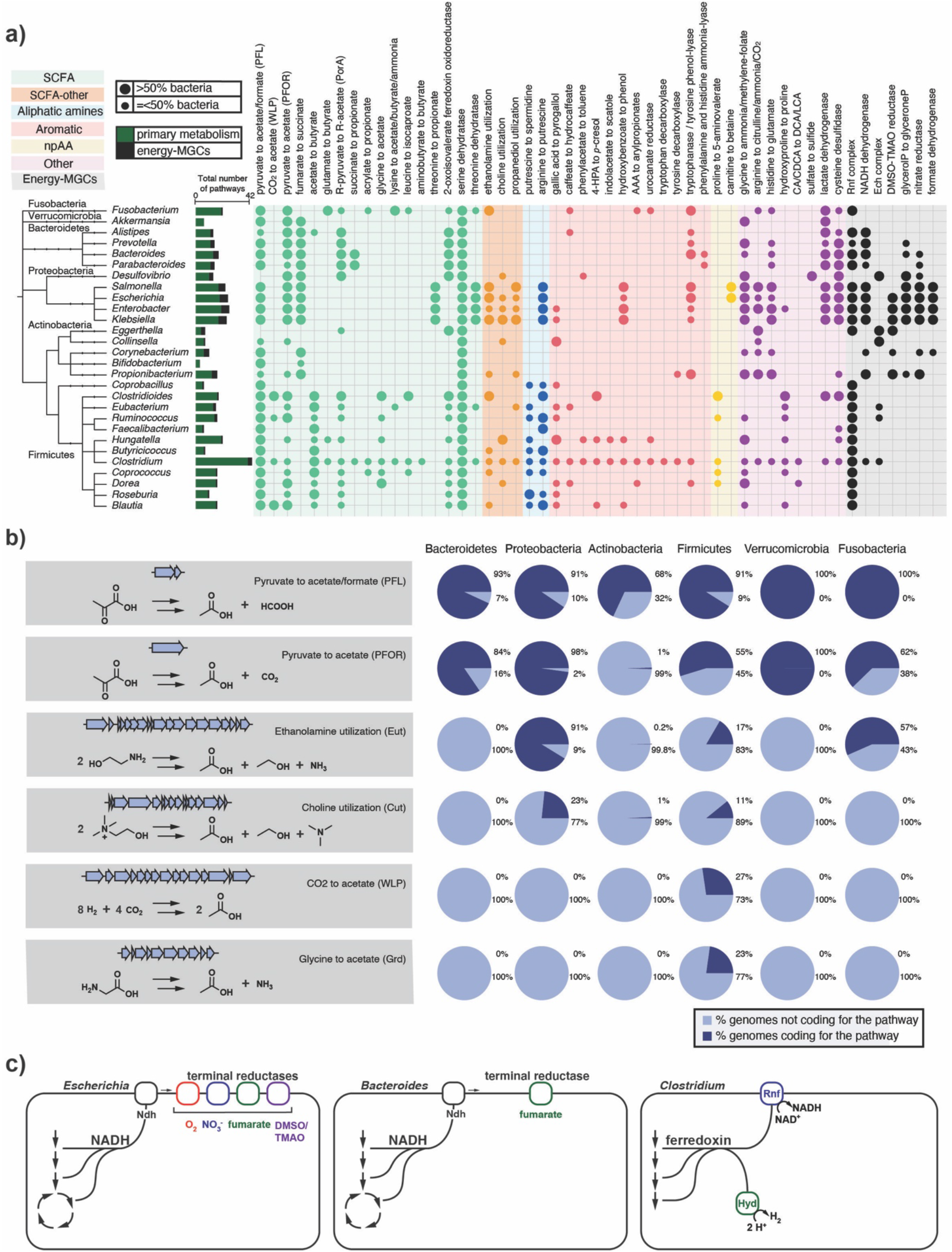
Distribution of known pathways across most representative genera in the human gut. (A) Circles represent the absence/presence of known pathways in each genus. Larger circles indicate cases in which more than 50% of the genomes for a genus encode the pathway, while smaller circles indicate cases in which 50% or fewer of the genomes encode it. Colored ranges indicate a categorization of MGCs by chemical class of their product, in which npAA represents nonproteinogenic amino acids and SCFA represents short-chain fatty acids. Taxonomic assignments were applied using the Genome Taxonomy Database (GTDB) (*10*). The tree was generated using phyloT (https://phylot.biobyte.de/) and visualized using iTOL (*11*). Raw data are available in Table S5. (B) Distribution of the main acetate synthesis pathways at phylum level. Some of the pathways are ubiquitous across the five major phyla (e.g. pyruvate to acetate/formate [PFL]), while others are only found in Firmicutes (CO2 to acetate [WLP]). Raw data for the pie charts is available in Table S6. Genes and gene clusters depicted are representatives from *Bacteroides thetaiotaomicron* (PFL & PFOR), *Salmonella enterica* (Eut), *Clostridium sporogenes* (Cut), *Clostridium difficile* (WLP) and *Clostridium sticklandii* (Grd). (C) Bioenergetic strategies in *Escherichia* that has a variety of alternate electron acceptors to choose from compared to *Bacteroides* and *Clostridium*. Abbreviations: PFL, pyruvate formate-lyase; PFOR, pyruvate:ferredoxin oxidoreductase; Eut, ethanolamine utilization; Cut, choline utilization; WLP, Wood-Ljungdahl Pathway; Grd, glycine reductase; CA, cholic acid; CDCA, chenodeoxycholic acid; DCA, deoxycholic acid; LCA, lithocholic acid; TMAO, trimethylamine N-oxide; DMSO, dimethylsulfoxide; SCFA, short-chain fatty acid; Ndh, NADH dehydrogenase, Rnf, Rhodobacter nitrogen fixation like complex; Hyd, hydrogenase.

Our results provide insights into the metabolic strategies that microbes use to produce short chain fatty acids (SCFAs). As expected, butyrate production is found exclusively in certain Firmicutes and Fusobacteria, whereas propionate production is largely confined to (and conserved in) the Bacteroidetes. However, the phylogenetic distribution of pathways that generate acetate -- the most concentrated molecule produced in the gut (*12*) -- has not yet been described. Two pathways for the conversion of pyruvate to acetate -- pyruvate formate-lyase (pyruvate to acetate/formate) and pyruvate:ferredoxin oxidoreductase (PFOR) -- are widely distributed across microbial strains from diverse phyla (Figure 2B). Two observations suggest that these two pathways are the most prolific source of acetate in the gut. First, some strains known to produce large quantities of acetate rely entirely on one or both of the pathways. Second, each one uses pyruvate as a substrate, consistent with a model in which these pathways are the primary conduit through which carbohydrate-derived carbon is converted to acetate. Additional taxon-specific pathways for acetate include the CO_2_ to acetate pathway and the glycine to acetate pathway (each specific to a subset of Firmicutes), as well as the choline and ethanolamine utilization pathways (widespread among Enterobacteriaceae and each found in different clades of Firmicutes) (Figure 2A).

Our results demonstrate a striking difference in mechanisms for energy capture by three of the major bacterial genera in the gut: *Bacteroides, Escherichia*, and *Clostridium*. When growing aerobically with glucose, *E. coli* generates most of its energy by channelling electrons through membrane bound cytochromes using oxygen as the terminal electron acceptor (Figure 2C). However, oxygen is limiting in the gut. Under anaerobic conditions, bacteria from the genus *Escherichia* employ alternate terminal electron acceptors such as nitrate, DMSO, TMAO, and fumarate by substituting alternate terminal reductases into their electron transport system (Figure 2C). However, in the healthy gut these alternate electron acceptors are either absent or available in limited amounts, likely explaining why these facultative anaerobes represent a small proportion of the healthy microbiome (*13*). In contrast to the diversity of terminal reductases used by the *Escherichia, Bacteroides* genomes encode only fumarate reductase (Figure 2C). They use a unique pathway, carboxylating PEP to form fumarate, which they use as a terminal electron acceptor to run an anaerobic electron transport chain involving NADH dehydrogenase and fumarate reductase, ultimately forming propionate. Thus, the metabolic strategy employed by *Bacteroides* ensures a steady stream of electron acceptor to fuel their metabolism. The *Clostridium* do not utilize similar mechanisms for energy capture as the *Escherichia* and the *Bacteroides*. Recent analyses suggest that they use the Rnf complex for generating a proton motive force. Several pathways encoded by the genomes of *Clostridium* (e.g., acetate to butyrate, AAA to arylpropionates, leucine to isocaproate) (Figure 2A) consist of an electron bifurcating acyl-CoA dehydrogenase enzyme. This complex bifurcates electrons from NADH to the low potential electron carrier ferredoxin which can then donate electrons to the RNF complex which functions as a proton or sodium pump, generating an ion motive force. Although much still is to be learned about Clostridial metabolism, our findings suggest that their metabolism operates at a different scale of the redox tower compared to *Bacteroides* and Enterobacteriaceae, using low potential electron carriers to fuel their metabolism.

Next, we set out to determine the prevalence and abundance of each pathway in a cohort of human samples. We used BiG-MAP (*14*) to profile the relative abundance of each MGC class across 1,135 metagenomes from the population-based LifeLines DEEP cohort (*15*), by mapping metagenomic reads against a collection of 6,836 non-redundant MGCs detected in our set of reference genomes (Figure 3A,B). Some pathways, such as CO_2_ to acetate (acetogenesis) and butyrate production from acetate or glutamate, as well as polyamine-forming pathways, were found in >99% of microbiomes. Others, such as 1,2-propanediol utilization and *p*-cresol production, both associated with negative effects on gut health (*16, 17*), were observed at detectable levels in only half of the samples. In terms of abundance, it is striking that for example the bile acid-induced (*bai*) operon for the formation of the secondary bile acids deoxycholic acid and lithocholic acid, which has been characterized from very low-abundance *Clostridium scindens* strains (*18*), was still shown to be present in relatively high abundance across a subset of subjects. Analysis of the mapped reads showed that the vast majority of these mapped to a homologous MGC from the genus *Dorea* instead (Suppl. Figure 2), for which the physiological relevance remains to be established. It is also interesting to see that, while two of the three acetate-forming pathways (PFL and WLP) were consistently found at high abundance levels, the abundance of all butyrate-forming pathways is highly variable across subjects, with a >20-fold difference between lower and upper quartiles in the abundance distribution of the glutamate-to-butyrate pathway, and a >440-fold difference between the 10th percentile and the 90th percentile.

**Figure 3.**
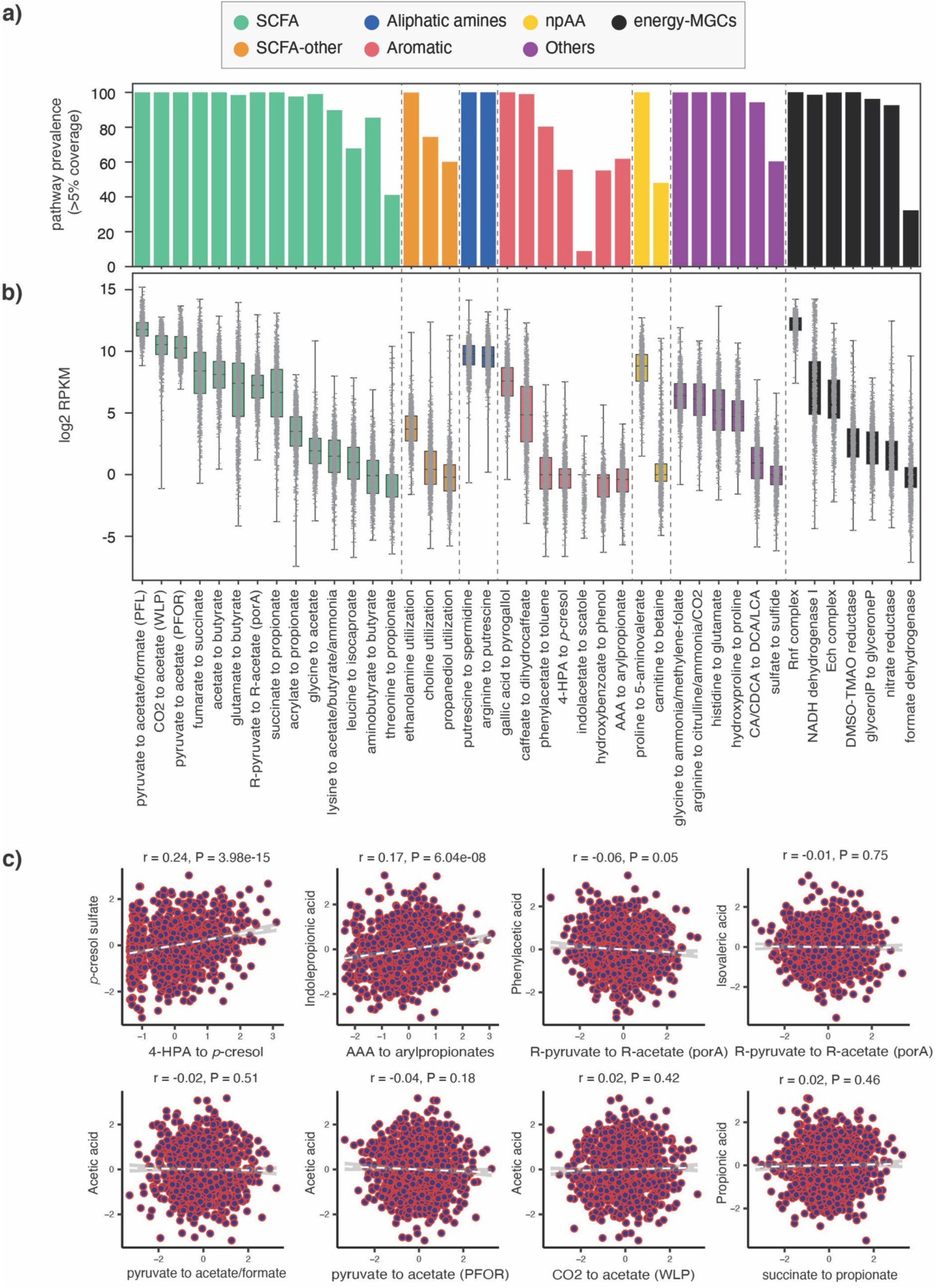
Prevalence and abundance of specialized primary metabolic pathways across 1,135 human microbiome samples. (A) Prevalence of each of the 41 known pathway classes across all microbiomes, measured as the percentage of samples in which core enzyme-coding genes of at least one reference MGC belonging to a given class were covered by metagenomic reads across >5% of their sequence length. This cutoff was kept low to avoid false negatives due to limited sequencing depth for low-abundance taxa (raw data available at Table S8). (B) Distributions of log2 RPKM relative abundance values of all 41 known pathway classes, categorized by product class, across all LifeLines DEEP metagenomes (raw data available at Table S9). (C) Limited correlation of genetic pathway abundance with abundance of metabolites in blood plasma.

The wide variability in the metagenome abundance of each pathway raises the question of whether metagenomic abundance of a pathway correlates with the level of its small molecule product in the host. To address this question, we systematically compared the level of each pathway with the quantity of the corresponding metabolite as determined by plasma metabolomics. We find a striking lack of correlation between pathway and metabolite levels (*r* ranging from −0.04 to 0.24, Figure 3C). These data indicate that gene abundances in metagenomes are not (on their own) a useful predictor of metabolic outputs. This finding has important implications for analyses that make metabolic inferences from gene abundances (*19*) or the abundances of individual strains (*20*). We speculate that a more detailed understanding of the influence of diet, differences in gene regulation, characteristic pathway flux (turnovers per unit time per protein copy), and pharmacokinetic characteristics (e.g., absorption, distribution, metabolism, and excretion) could ultimately enable the prediction of metabolite abundance from metagenome abundance. The systematic detection of the relevant genes and gene clusters by gutSMASH will provide a technological foundation for future studies in this direction, by allowing mapping of metatranscriptomic data to these accurately defined and categorized sets of genomic loci in order to understand which conditions and interactions are driving the expression of these pathways and the accumulation of their products.

The gutSMASH software constitutes, to our knowledge, the first comprehensive automated tool designed to identify niche-defining primary metabolic pathways from genome sequences or metagenomic contigs—even a full-fledged metabolic network reconstruction software like PathwayTools (*21*) (which uses the extensive MetaCyc database (*22*)) lacks detection capabilities for 3 out of the 41 MGC-encoded pathways detected by gutSMASH (Table S7). Moreover, the identification of MGCs provides considerably increased confidence that detected homologues for a given pathway are truly working together. Downstream, detected MGCs can be used as input for read-based tools such as HUMAnN (*23*) or BiG-MAP (*14*) to measure abundance or expression levels of the encoded pathways. On top of these functionalities, the gutSMASH framework also facilitates identifying new (i.e., uncharacterized) pathways in the microbiome. To this end, we designed an additional set of rules to detect primary metabolic gene clusters of unknown function that harbor Fe-S flavoenzymes (*24*), glycyl-radical enzymes, 2-hydroxyglutaryl-CoA-dehydratase-related enzymes, and/or enzymes involved in oxidative decarboxylation. From this analysis of putative MGCs (see SI methods *Analysis of distant homologues and putative MGCs from CGR, HMP and Clostridioides dataset*), we found 12,259 putative MGCs from 760 different species, that, after redundancy filtering at 90% sequence similarity, were classified into 932 GCFs. Within these, we manually prioritized a range of gene clusters with unprecedented enzyme-coding gene content (see Suppl. Figure 4&5, SI Results *Analysis of putative clusters and distant homologues: relevant candidates to study further*).These can be a potential new source to discover new pathways and metabolites.

## References

1. K. Blin, S. Shaw, K. Steinke, R. Villebro, N. Ziemert, S. Y. Lee, M. H. Medema, T. Weber, antiSMASH 5.0: updates to the secondary metabolite genome mining pipeline. Nucleic Acids Res. 47, W81–W87 (2019).

2. P. D. Karp, R. Billington, R. Caspi, C. A. Fulcher, M. Latendresse, A. Kothari, I. M. Keseler, M. Krummenacker, P. E. Midford, Q. Ong, W. K. Ong, S. M. Paley, P. Subhraveti, The BioCyc collection of microbial genomes and metabolic pathways. Brief. Bioinform. 20, 1085–1093 (2019).

3. S. Abubucker, N. Segata, J. Goll, A. M. Schubert, J. Izard, B. L. Cantarel, B. Rodriguez-Mueller, J. Zucker, M. Thiagarajan, B. Henrissat, O. White, S. T. Kelley, B. Methé, P. D. Schloss, D. Gevers, M. Mitreva, C. Huttenhower, Metabolic reconstruction for metagenomic data and its application to the human microbiome. PLoS Comput. Biol. 8, e1002358 (2012).

4. J. C. Navarro-Muñoz, N. Selem-Mojica, M. W. Mullowney, S. Kautsar, J. H. Tryon, E. I. Parkinson, E. L. C. De Los Santos, M. Yeong, P. Cruz-Morales, S. Abubucker, A. Roeters, W. Lokhorst, A. Fernandez-Guerra, L. T. D. Cappelini, R. J. Thomson, W. W. Metcalf, N. L. Kelleher, F. Barona-Gomez, M. H. Medema, A computational framework for systematic exploration of biosynthetic diversity from large-scale genomic data. Nat. Chem. Biol. 16, 60–68 (2020).

5. S. Kitamoto, C. J. Alteri, M. Rodrigues, H. Nagao-Kitamoto, K. Sugihara, S. D. Himpsl, M. Bazzi, M. Miyoshi, T. Nishioka, A. Hayashi, T. L. Morhardt, P. Kuffa, H. Grasberger, M. El-Zaatari, S. Bishu, C. Ishii, A. Hirayama, K. A. Eaton, B. Dogan, K. W. Simpson, N. Inohara, H. L. T. Mobley, J. Y. Kao, S. Fukuda, N. Barnich, N. Kamada, Dietary L-serine confers a competitive fitness advantage to Enterobacteriaceae in the inflamed gut. Nat. Microbiol. 5, 116–125 (2020).

6. Y. Zou, W. Xue, G. Luo, Z. Deng, P. Qin, R. Guo, H. Sun, Y. Xia, S. Liang, Y. Dai, D. Wan, R. Jiang, L. Su, Q. Feng, Z. Jie, T. Guo, Z. Xia, C. Liu, J. Yu, Y. Lin, S. Tang, G. Huo, X. Xu, Y. Hou, X. Liu, J. Wang, H. Yang, K. Kristiansen, J. Li, H. Jia, L. Xiao, 1,520 reference genomes from cultivated human gut bacteria enable functional microbiome analyses. Nat. Biotechnol. 37, 179–185 (2019).

7. J. Lloyd-Price, A. Mahurkar, G. Rahnavard, J. Crabtree, J. Orvis, A. B. Hall, A. Brady, H. H. Creasy, C. McCracken, M. G. Giglio, D. McDonald, E. A. Franzosa, R. Knight, O. White, C. Huttenhower, Strains, functions and dynamics in the expanded Human Microbiome Project. Nature 550, 61–66 (2017).

8. B. P. Tracy, S. W. Jones, A. G. Fast, D. C. Indurthi, E. T. Papoutsakis, Clostridia: the importance of their exceptional substrate and metabolite diversity for biofuel and biorefinery applications. Curr.Opin. Biotechnol. 23, 364–381 (2012).

9. F. Maguire, B. Jia, K. L. Gray, W. Y. V. Lau, R. G. Beiko, F. S. L. Brinkman, Metagenome-assembled genome binning methods with short reads disproportionately fail for plasmids and genomic Islands. Microb. Genom. 6, mgen.0.000436 (2020).

10. D. H. Parks, M. Chuvochina, D. W. Waite, C. Rinke, A. Skarshewski, P.-A. Chaumeil, P. Hugenholtz, A standardized bacterial taxonomy based on genome phylogeny substantially revises the tree of life. Nat. Biotechnol. 36, 996–1004 (2018).

11. I. Letunic, P. Bork, Interactive Tree Of Life (iTOL) v4: recent updates and new developments. Nucleic Acids Res. 47, W256–W259 (2019).

12. J. H. Cummings, E. W. Pomare, W. J. Branch, C. P. Naylor, G. T. Macfarlane, Short chain fatty acids in human large intestine, portal, hepatic and venous blood. Gut 28, 1221–1227 (1987).

13. S. A. Jones, T. Gibson, R. C. Maltby, F. Z. Chowdhury, V. Stewart, P. S. Cohen, T. Conway, Anaerobic respiration of Escherichia coli in the mouse intestine. Infect. Immun. 79, 4218–4226 (2011).

14. V. P. Andreu, H. E. Augustijn, K. van den Berg, J. J. J. van der Hooft, M. A. Fischbach, M. H. Medema, BiG-MAP: an automated pipeline to profile metabolic gene cluster abundance and expression in microbiomes. BioRxiv, doi:10.1101/2020.12.14.422671 (2020).

15. E. F. Tigchelaar, A. Zhernakova, J. A. M. Dekens, G. Hermes, A. Baranska, Z. Mujagic, M. A. Swertz, A. M. Muñoz, P. Deelen, M. C. Cénit, L. Franke, S. Scholtens, R. P. Stolk, C. Wijmenga, E. J. M. Feskens, Cohort profile: LifeLines DEEP, a prospective, general population cohort study in the northern Netherlands: study design and baseline characteristics. BMJ Open 5, e006772 (2015).

16. F. Faber, P. Thiennimitr, L. Spiga, M. X. Byndloss, Y. Litvak, S. Lawhon, H. L. Andrews-Polymenis, S. E. Winter, A. J. Bäumler, Respiration of Microbiota-Derived 1,2-propanediol Drives Salmonella Expansion during Colitis. PLoS Pathog. 13, e1006129 (2017).

17. M. Andriamihaja, A. Lan, M. Beaumont, M. Audebert, X. Wong, K. Yamada, Y. Yin, D. Tomé, C. Carrasco-Pozo, M. Gotteland, X. Kong, F. Blachier, The deleterious metabolic and genotoxic effects of the bacterial metabolite p-cresol on colonic epithelial cells. Free Radic. Biol. Med. 85, 219–227 (2015).

18. M. Funabashi, T. L. Grove, M. Wang, Y. Varma, M. E. McFadden, L. C. Brown, C. Guo, S. Higginbottom, S. C. Almo, M. A. Fischbach, A metabolic pathway for bile acid dehydroxylation by the gut microbiome. Nature 582, 566–570 (2020).

19. H. Mallick, E. A. Franzosa, L. J. Mclver, S. Banerjee, A. Sirota-Madi, A. D. Kostic, C. B. Clish, H. Vlamakis, R. J. Xavier, C. Huttenhower, Predictive metabolomic profiling of microbial communities using amplicon or metagenomic sequences. Nat. Commun. 10, 3136 (2019).

20. G. M. Douglas, V. J. Maffei, J. R. Zaneveld, S. N. Yurgel, J. R. Brown, C. M. Taylor, C. Huttenhower, M. G. I. Langille, PICRUSt2 for prediction of metagenome functions. Nat. Biotechnol. 38, 685–688 (2020).

21. P. D. Karp, P. E. Midford, R. Billington, A. Kothari, M. Krummenacker, M. Latendresse, W. K. Ong, P. Subhraveti, R. Caspi, C. Fulcher, I. M. Keseler, S. M. Paley, Pathway Tools version 23.0 update: software for pathway/genome informatics and systems biology. Brief. Bioinform. 22, 109–126 (2021).

22. R. Caspi, R. Billington, I. M. Keseler, A. Kothari, M. Krummenacker, P. E. Midford, W. K. Ong, S. Paley, P. Subhraveti, P. D. Karp, The MetaCyc database of metabolic pathways and enzymes - a 2019 update. Nucleic Acids Res. 48, D445–D453 (2020).

23. E. A. Franzosa, L. J. McIver, G. Rahnavard, L. R. Thompson, M. Schirmer, G. Weingart, K. S. Lipson, R. Knight, J. G. Caporaso, N. Segata, C. Huttenhower, Species-level functional profiling of metagenomes and metatranscriptomes. Nat. Methods. 15, 962–968 (2018).

24. V. Pascal Andreu, M. A. Fischbach, M. H. Medema, Computational genomic discovery of diverse gene clusters harbouring Fe-S flavoenzymes in anaerobic gut microbiota. Microb. Genom. 6, mgen.0.000373 (2020).

