## Supplementary material for "A systematic analysis of metabolic pathways in the human gut microbiota"

### Supplementary material for:
### A systematic analysis of metabolic pathways in the human gut microbiota

Victòria Pascal Andreu[1], Hannah E. Augustijn[1,2#], Lianmin Chen[2,3,#], Alexandra Zhernakova[2], Jingyuan Fu[2,3], Michael A. Fischbach[4,5,7*], Dylan Dodd[5,6*], Marnix H. Medema[1*]

#### Supplementary Results

##### *Phylogenetic analysis of protein superfamilies to identify pathway-specific clades*

Performing hmmscan searches on the protein sequences helped identify the presence of keystone domains (Pfams) between pathways that share an enzymatic core. However, some of these enzymes are part of multifunctional enzyme families that are recognised by very broad Pfam domains. Thus, in order to more accurately identify relevant functional subgroups for keystone enzyme families, we used protein phylogenetic analysis to pinpoint pathway-specific clades and discern, for instance, specialized primary metabolism-related enzymes from housekeeping related ones (Figure 1). For each protein family, we created a non-redundant set of representatives by gathering protein sequences from four different sources: the proteins from the representative pathway, the reference proteome available in Pfam (*1*), the respective proteins from the MGC collection and the experimentally characterized proteins available in UniProt, to assign functionality to clades (see SI Methods: *Towards a more robust MGC identification by building new HMM profiles*). In total, we performed this phylogenetic analysis to 12 major protein superfamilies. For instance, phenyllactate dehydratase homologues are involved in the degradation of aromatic amino acids into propionate, in the acrylate to propionate pathway and in the leucine reductive branch pathway; in contrast, 2-hydroxyglutaryl-CoA dehydratase allows the transformation of glutamate into butyrate. Despite the fact that these reactions are different, both key enzymes harbour the 2-hydroxyglutaryl-CoA dehydratase, D-component domain (HGD-D Pfam domain, PF06050). The HGD-D protein family phylogenetic tree (see Suppl. Figure 3) revealed that 10 clades were implicated in these three pathways. Subsequently, 10 profile hidden Markov models (pHMMs) specific to these clades were built and, after assessing the sensitivity of the models (see Methods), 9 out of 10 were selected for being sensitive enough as to correctly identify the subdomains of this protein family involved in these 3 pathways. Consequently, these 9 pHMMs were included in the corresponding detection rule. The same procedure was followed for the other protein superfamilies, creating a total of 43 pHMMs (see Table S10). As a result, gutSMASH uses newly built pHMMs in combination with the ones included in the Pfam database to identify proteins families of interest. Therefore, gutSMASH competitively scores hmmsearch hits and assigns to the sequence the domain with a higher score. Altogether, this procedure allowed to define a preliminary set of detection rules using the newly built pHMMs to predict close homologues of known gene clusters.

##### *Validation of gutSMASH detection rules by evaluating their predictive performance*

Evaluating gutSMASH performance implies having a set of bacteria whose genomes are known to encode a given pathway. However, the false positive rate is unknown (as it is not feasible to experimentally verify large numbers of diverse putative annotations) and the false negative rate is difficult to determine (as only in few species, MGCs with proven functions are described in literature).

Hence, we decided that the best course of action would be to perform detailed manual analysis of large numbers of diverse predicted MGCs. For this reason, gutSMASH was run on a test set created from 1,632 bacterial genomes. All the MGCs predicted by the same detection rule were grouped together to further run BiG-SCAPE on each subset (see Methods section *Testing and validating gutSMASH specific-to-known-pathway detection rules*). In this manner, we could evaluate the range of gene clusters predicted by the same detection rule and quickly find out if any distantly-related gene cluster should not be picked by the rule based on, e.g., having divergent enzyme-coding composition. Also, it allowed us to acquire an overview of the bacterial taxa predicted to possess a given gene cluster type and identify if any MGCs from taxa referenced in primary literature were missing from this set. Thus, this system allowed to fine-tune the detection rules and evaluate their predicting potential. After several iterations of adjusting rules, performing new predictions and creating new sequence similarity networks, we froze gutSMASH version 1.0 with 41 specific-to-known-pathway detection rules (see Table S11) to accurately and comprehensively predict MGCs.

An additional validation step was performed using PaperBLAST (*2*), which was used to look for genomes encoding any close homologues of the key proteins involved in 18 gutSMASH predicted pathways; all 18 MGCs were successfully detected using gutSMASH. When compared to the reference MGCs, detected clusters showed an average amino acid sequence identity between 54,46 and 100% and overall gene cluster similarity (percentage of homologues detected in KnownClusterBlast) ranging from 44 to 100%.

***Analysis of putative clusters and distant homologues: relevant candidates to study further***

Next, we evaluated the potential of gutSMASH to predict putative MGCs of interest and to explore the metabolic landscape covered by them. For this reason, the 12,259 putative clusters predicted from the HMP, CGR and *Clostridioides* genomes were used and subjected to a redundancy filtering of 90% similarity at the protein sequence level (see Methods: *Evaluating the functional potential of the human microbiome using gutSMASH*) to investigate their functional diversity. To create a non-redundant collection, two random representative clusters of each set of highly similar clusters were picked; all representatives were then clustered together into 932 GCFs using BiG-SCAPE (*3*) (see Methods: *Analysis of distant homologues and putative MGCs from CGR, HMP and Clostridioides dataset*). From the resulting network (see Suppl. Figure 4), we made three main observations. First, we identified several distant homologues of the known MGCs; these are picked by specific pathway rules but classified as putative when for instance the MGC is from a distantly related taxonomic group and therefore shares low sequence similarity with the reference gene cluster. Second, we found previously characterized MGCs that were not included in the original training sets of known MGC types, as for instance a region from *Ruminococcus gnavus strain AM22-7AC* (accession number QRIA01000012.1) involved in the metabolism of rhamnose and fucose described in 2013 by E.Petit & W. Latouf *et al* [ref]; this validated the capability of gutSMASH to identify real MGCs that were not included in the initial list of known pathways. Third, we observed vast numbers of novel clusters of unknown function that represent good candidates for further experimental characterization. The examples shown in Suppl. Figure 4 are metabolically diverse MGCs that encode flavoenzymes, oxidative decarboxylation (OD), glycyl radical (GR), 2-hydroxyglutaryl-CoA dehydratase (HGD-D) related or hybrids of them, which were found in phylogenetically diverse genomes from Firmicutes, Fusobacteria and Actinobacteria. Moreover, all these MGCs presented plausible architectures as to be real gene clusters, since they included all the genetic elements to regulate, synthesize and transport the resulting molecule (or its substrates). The first MGC for instance, which is found in *Enterocloster* *citronae* AF29-4, shares some similarity with the acetate to butyrate MGCs, since it encodes an acyl-CoA dehydrogenase and two electron transfer flavoproteins as well as a distant *baiH* homologue. Another interesting example is the *Dorea sp.* D27 MGC, a pathway involving a pyruvate:ferredoxin oxidoreductase, which might be encapsulated given the presence of bacterial microcompartment genes (BMC-encoding genes) in the cluster. *Blautia sp.* TF11-31AT also presents an unprecedented gene cluster architecture, encoding a combination of enzymes involved in oxidative decarboxylation (thiamine pyrophosphate-related) and related to HGD-D. This overview highlights the ability of gutSMASH to systematically predict novel and interesting gene clusters from a diverse range of bacteria that could help associating function to unknown genes and predict novel pathways. Additionally, the expression of families of MGCs of unknown function could potentially be correlated to microbiome-associated phenotypes to prioritize them for experimental characterization based on physiological and ecological relevance.

***Assessing pathway abundance and prevalence across metagenomes***

From the predicted pool of known pathways, we aimed to assess their abundance and prevalence by mapping the metagenomic reads of the LifeLines-DEEP cohort (*4*, *5*). To compute both, certain assumptions had to be made and thresholds needed to be chosen. In both cases, the numbers of reads mapping to the core regions are key to assess whether a given pathway is present or not and how abundant it is. In order to account for spurious mapping, we designed an approach to assess the pathway abundance by using the lower quartile number of reads mapping to 2kb long regions for each pathway and sample (see more in Methods section entitled *Mapping metagenomics reads from healthy samples to the known gutSMASH predicted MGCs*). In contrast, to evaluate the prevalence of all the pathways across samples, the core coverage score estimated by BiG-MAP was used. Ideally, with unlimited sequencing depth, one would expect to find reads mapping evenly to the whole gene cluster, thus, implying a relatively high core coverage score compared to the chosen one (>5% coverage, see Figure 5) and also account for spurious mapping. However, when raising the minimum coverage value from 10-80% some of the pathways that are known to be present in healthy individuals showed a prevalence of 0 (see Suppl. Figure 6). Assessing the minimum sequence identity of a read mapping to a gene cluster showed 78% similarity (at nucleotide level), which confirms that even with low coverage scores, the reads are mapping very specifically to the gene clusters and hence, the lack of coverage of some gene clusters is very likely due to the sequence depth of a sample rather than the absence of the pathway. For this reason, we set the minimum coverage used in the analyses presented in the main text at 5%, to avoid undue false negatives.

#### Methods

gutSMASH is a Python-based pipeline that has been built from antiSMASH version 5.0 source code. The latest command line version is freely available and can be downloaded and installed from here: https://github.com/victoriapascal/gutsmash/tree/gutsmash

##### *Finding pathway signatures for a collection of known and characterized MGCs*

To create a new set of detection rules, 41 known and characterized MGCs were gathered from literature and used as positive controls. The protein sequences of these MGCs were searched using hmmscan (HMMER suit version 3.1b2, February 2015; http://hmmer.org/). From the resulting pHMM profile hits, auxiliary and core domains were manually identified for each pathway, to ultimately determine the pathway signature and specify it in the corresponding detection rule. To discern and more precisely identify key enzymes of interest sharing a keystone domain, we used custom-made pHMMs following a procedure described in the SI Methods section: *Towards a more robust MGC identification by building new HMM profiles*. Altogether, the knowledge on the core enzyme coding-genes and the newly-built pHMMs helped to construct a preliminary set of detection rules to predict known pathways.

##### *Towards a more robust MGC identification by building new HMM profiles*

Certain core domains are shared across diverse pathways, including the PFL-like domain and the HGD-D domain. In total, 13 keystone domains were found to be ubiquitous in multiple pathways (see SI Table 10). Hence, to increase gutSMASH precision and discern between enzyme subfamilies of interest, 12 protein superfamily phylogenies were constructed by aligning the protein sequences harbouring the domain of interest from the MGC collection (described in Methods section *Exploring the yet unknown metabolic diversity by creating general detection rules*), the respective reference proteome[34] at a 15% or 35% co-membership threshold (the latter only for the domains Gly_radical and Acyl-CoA_dh_1) and any experimentally characterized UniProt representatives. After aligning the sequences with Clustal Omega (*6*), approximately-maximum-likelihood phylogenetic trees using FastTree 2.1 (*7*) were inferred to further annotate the tree with iTOL (*8*). Thus, from the desired and functionally relevant clades, specific pHMMs were built by extracting the amino acid sequence of the clade-specific proteins, aligning them with Clustal Omega, trimming the edges of the multiple sequence alignment using Jalview (*9*), re-aligning all the sequences with Clustal Omega and finally building a pHMM using hmmbuild (HMMER suite version 3.1b2, February 2015; http://hmmer.org/). Subsequently, for all the newly created pHMMs, sensitivity was assessed using 10-fold jackknife cross-validation. Each clade was divided randomly into training and testing sets. The protein sequences from the training set were aligned using Clustal Omega and used to create a pHMM. Next, the protein sequences of the test set were hmmscanned (HMMER suit version 3.1b2, February 2015; http://hmmer.org/) against the newly built testing pHMMs. When a sequence scored positively for multiple domains in the same region, only the domain with a higher bit score was picked out. Sensitivity then accounted for the number of sequences positively associated with the correct pHMM out of the total number of sequences in the testing set. The same procedure was repeated 10 times. The pHMMs with a true positive rate higher than 0.85 across the 10 rounds were included in the detection rules. In total, 43 newly built pHMMs were included in the corresponding detection rules. Moreover, a pHMM to capture succinate dehydrogenase/fumarate reductase was built by aligning 10 protein sequences of such enzymes and building the model from this alignment using a hmmbuild. To also competitively score similar Pfam domains, HHsearch pre-computed results obtained from the Pfam FTP (ftp://ftp.ebi.ac.uk/pub/databases/Pfam/current_release/database_files/) were parsed and included in the gutSMASH code.

***Testing and validating gutSMASH specific-to-known-pathway detection rules***

To evaluate the performance of the preliminary set of detection rules, a total of 1,621 bacterial genomes, including 1,520 genomes from the CGR collection (*10*) and 101 manually selected genomes from the most representative bacterial genera in the human gut, were used as input for gutSMASH (see Table S3). The predicted MGCs were classified based on the detection rule they were predicted from, to later run BiG-SCAPE on each sub-collection. The resulting networks were screened individually to evaluate the taxonomic and architectural diversity, to assess if any architectural variant or taxon (based on literature) was missing from the MGC pool or was incorrectly predicted by the detection rule. Hence, this procedure ultimately helped to tweak the detection rules to predict true homologues of the known pathways. After two iterations of fine-tuning and testing, all detection rules were performing as intended and constituted the new set of detection rules of gutSMASH version 1.0.

***gutSMASH customized databases and output visualization***

The antiSMASH version 5.0 source code was further tailored to meet gutSMASH functionality. The 32,144 predicted MGCs obtained from running gutSMASH on the CGR, HMP and Clostridiales collections (see Methods section *Evaluating functional potential of gut bacteria using gutSMASH* for more insights), were used to create the CluterBlast database. In a similar way, 59 positive controls carrying the known pathways (from which we created the specific-to-known-pathway detection rules) were used to create the KnownClusterBlast database. These databases facilitate comparative gene cluster analysis using BLAST (*11*). Thus, they allow assessing how broadly distributed an MGC is across bacteria (in the case of ClusterBlast) or evaluating the similarity between the predicted MGC and a known and functionally characterized MGC (when using KnownClusterBlast).

Another functionality of antiSMASH is to classify coding genes based on the domains into five major functional categories: core biosynthetic, additional biosynthetic, transport-related, regulatory, resistance and other, using the pmCOG (primary metabolism Clusters of Orthologous Groups) tool, which is embedded in antiSMASH (there originally named smCOG for 'secondary' metabolism Clusters of Orthologous Groups). Thus, the pHMM library pmCOG uses was updated to include relevant domains found in specialized primary metabolism. Also, two other important functional categories were added: electron transport-related genes and encapsulation genes.

***Exploring the yet unknown metabolic diversity by creating general detection rules***

With the objective of creating general detection rules to predict putative MGCs, a similar approach used to screen the surrounding genes around a Fe-S flavoenzyme coding gene was used (*12*). Some of the representative known pathways share proteins with biochemically similar functions; these include, for instance, pyruvate formate lyase-like enzymes that are found in the threonine-to-propionate pathway, the choline utilization pathway and the pyruvate-to-acetate pathways. In order to cover a large amount of sequence diversity, we created a database that included 11,000 complete genomes and 98,886 draft genomes available in Genbank (in February 2017) in order to use clusterTools (*13*), a software to find remote homologues of known MGCs. As input, a subset of the known pathways used to design the detection rules for known pathways were used as input (see Table

S12). The output of several iterated clusterTools searches were grouped to acquire a collection of over 29,000 clusters. For visualization and manual scoring purposes, MultiGeneBlast (*14*) was run using the clusterTools output as input. Thus, MGCs harbouring at least half of the genes from the query gene list and with a cumulative BLAST score higher than 1,000 were included in the MGC collection. In order to filter out redundant sequences, we used MMseqs2 (*15*) at a 95% similarity cut-off. From the resulting network of 1,599 groups, a maximum of 1 random representative plus singletons were picked creating a 'non-redundant' set of almost 3,200 clusters. This collection was screened for gene clusters harbouring the *baiCD* or *baiH* coding gene (Oxidored_FMN and Pyr_redox_2), pyruvate-formate lyase (PFL-like or Gly_radical), pyruvate ferredoxin (POR, POR_N or PFOR_II), thiamine pyrophosphate enzyme (TPP_enzyme_C) and 2-hydroxyglutaryl-CoA dehydratase (HGD-D), each of which are keystone domains in charge of important anaerobic reactions. This helped creating general detection rules, by identifying which other enzyme-coding Pfam domains are found around these 'anchor' domains in flanking regions; this was systematically analyzed per gene cluster family to make sure that the general rules captured all major families of homologous MGCs of interest. Also, when validating the specific-to-known pathway detection rules, whenever a specific rule predicted interesting MGCs that were variants of the representative pathway with likely differing functions, a general rule was created out of the specific one by loosening up the Pfam requirements. The full list of general rules can be found in Table S13.

***Assessing single-protein pathway abundance within representative human gut bacteria***

To include single-protein pathways in our analysis to assess the overall abundance of specialized primary metabolic pathways, 10 enzyme families were selected for downstream analysis. Following the same procedure as described in the Methods section: *Towards a more robust MGC identification by building new HMM profiles*, protein phylogenies were built for each protein superfamily. Similarly, from the pathway-specific monophyletic clades, we built new pHMMs. A bitscore threshold for each newly built pHMM was calibrated in order to identify with high confidence proteins belonging to the same functional clades. To this end, the protein sequences that composed the superfamily phylogeny were subjected to an hmmsearch run with the new pHMM. The bitscore reported by hmmsearch for the most distantly related protein within the pathway-specific clade was chosen as the threshold for that specific pHMM. Next, the protein sequences from the CGR, HMP and Clostridiales collections (further information in Methods section: *Evaluating the functional potential of the human microbiome using gutSMASH)* were scanned using the newly built pHMMs. Finally, the hmmsearch output tables for each pHMM were parsed so that the proteins with a bitscore equal or higher to the chosen threshold were deemed hits. In those cases in which the single-protein sequence codes for two Pfam domains, as for instance the serine dehydratase (SDH_alpha and SDH_beta), one of the Pfam domains was selected to create a protein phylogeny to further build a clade-specific pHMM, in this case SDH_alpha. Then, the protein sequences from the three collections were subjected to hmmsearch runs with both the clade-specific pHMM and the other co-occurring Pfam domain (in this case SDH_beta). The sequences that harbour both the specific pHMM at the chosen threshold and the co-occurring domain with an e-value =< $10^{-05}$ were deemed hits.

***Evaluating the functional potential of the human microbiome using gutSMASH***

To evaluate the metabolic potential of the human microbiome, gutSMASH was run on three different genome collections: (1) the CGR collection, with 1,520 CGR genomes deposited under the PRJNA482748, (2) the HMP reference genomes, with 2,146 HMP bacterial genomes downloaded in

September 2019 from here: https://www.hmpdacc.org/hmp/catalog/grid.php?dataset=genomic and (3) 414 Clostridiales complete genomes under the taxid 186802. The genomic FASTA sequence of these genomes was used as input for gutSMASH, which used Prodigal (*16*) to annotate genes across all of them in a consistent way. Moreover, in order to assess which MGC belonged to known pathways, the KnownClusterBlast (see SI Methods *gutSMASH customized databases and output visualization*) option was enabled. Thus, from the KnownClusterBlast output, the predicted regions were classified as known when the following two requirements were met: (1) an overall pathway similarity of at least 50% and at least half of the genes with a minimum protein sequence similarity of 40% or (2) an overall similarity of 60% and half of the genes with protein sequence similarity higher than 30%. However, in order not to penalize MGCs with similar domain profiles but substantially larger sizes, the requirements to be considered "known" slightly changed for the KnownClusterBlast MGCs longer than 17 coding genes. In this case, the same requirements as described above were used but instead of considering candidates with at least half of the coding genes having either 30 or 40% minimum sequence identity, one third of the genes were required to be present with the same minimum sequence identity. This was the case for the ethanolamine utilization operon, the *bai* operon characterized from *C. scindens* ATCC35704 (CA/CDCA to DCA/LCA pathway), the acetyl-CoA pathway (CO2 to acetate (WLP)), the tetrathionate to thiosulfate pathway and the NADH dehydrogenase I complex. Thus, all the MGCs that did not satisfy these conditions were classified as putative MGCs. The phylogenetic tree in Figure 2 was generated using phyloT v2 (https://phylot.biobyte.de/). The GDTB database (*17*) was used to assign the taxonomy to the genomes of the three collections (when present) and those taxonomic identifiers were the ones used for the subsequent pathway absence/presence analysis. Finally, the tree was annotated using iTOL (*8*).

***Analysis of distant homologues and putative MGCs from CGR, HMP and Clostridioides dataset***

The putative MGCs predicted from the CGR, HMP and Clostridiales genome collections were selected following the definition of "known" and "putative" gene clusters stated in the *Evaluating the functional potential of the human microbiome using gutSMASH* Methods section. To account for redundant MGCs, protein sequences extracted from all gene clusters were subjected to a redundancy filtering of 90% sequence similarity using MMseqs2. From the resulting clustering, two random representatives were chosen from each group, including the singletons. The resulting non-redundant collection of 3,040 putative MGCs was used as input for BiG-SCAPE using the default thresholds. The network in Suppl. Figure 4 was constructed and annotated using Cytoscape (*18*).

***Mapping metagenomics reads from healthy samples to the known gutSMASH predicted MGCs***

The HMP, CGR and Clostridiales-predicted MGCs were used as input for BiG-MAP (*19*), a tool that assesses gene cluster abundance or expression across metagenomics or metatranscriptomics data, respectively, by mapping the genomic reads onto the gene cluster sequences. The BiG-MAP family module grouped the 32,144 MGCs into 6,836 GCFs. Next, the reads of 1,135 participants of the population-based cohort LifeLines-DEEP (*5*) were mapped onto the resulting 6,836 Mash (*20*) representative MGCs by using BiG-MAP.map module. To assess the abundance of known pathways, the RPKM values from the known MGCs (following the definition of "known" stated in Methods section *Evaluating the functional potential of the human microbiome using gutSMASH*) were pulled out. The RPKM values of all the MGCs predicted by the same detection rule were merged. The pathway abundance (RPKM) was computed by dividing the gene clusters in 2kb-sized bins, and assessing the lower quartile number of reads mapping the 2kb bins for each gene cluster and sample. In contrast, a pathway was annotated as present in a sample when reads from that sample were found to be mapping to at least 5% of the core region of that MGC. This threshold was kept low to enable detection of MGCs from low-abundant microbes and avoid false negatives due to limited sequencing depth. The lowest percentage identity of reads mapped to MGCs was 78% at the nucleotide level, which instilled confidence that finding multiple reads mapping to different locations within a MGC provides sufficient evidence for its presence in a sample. The pathway prevalence was also computed using 10%, 20%, 30%, 40%, 50%, 60%, 70% and 80% core coverage thresholds (Suppl. Figure 5), and results for increasing thresholds were consistent with gradual loss of detection capability for pathways known to be associated with low-abundance bacteria, such as the AAA to arylpropionate pathway (aromatic amino acid reductive branch).

***Correlating pathway abundance with metabolite concentrations in plasma***

To evaluate the correlation between the gene cluster abundance and metabolite concentrations, the masses of 7 metabolites derived from several gutSMASH predicted gene clusters could be found in the Mass Spectrometry (MS) data of the plasma measured in LifeLines DEEP (*4*, *5*). These metabolites included acetic acid, indolepropionic acid, isovaleric acid, *p*-cresol, *p*-cresol sulfate, phenylacetic acid and propionic acid (see Figure 3c and Suppl. Figure 7). Both metabolite and pathway abundance (RPKM counts) were inverse-rank-transformed and the linear regression was applied to adjust covariates including age, sex and metagenomic sequencing depth (only for pathway abundance). Metabolite and pathway abundance residuals from the linear regression model were then used to perform Spearman correlation test. Finally, the Benjamini Hochberg method was applied to control for false discovery rate (FDR).

### Supplementary Figures

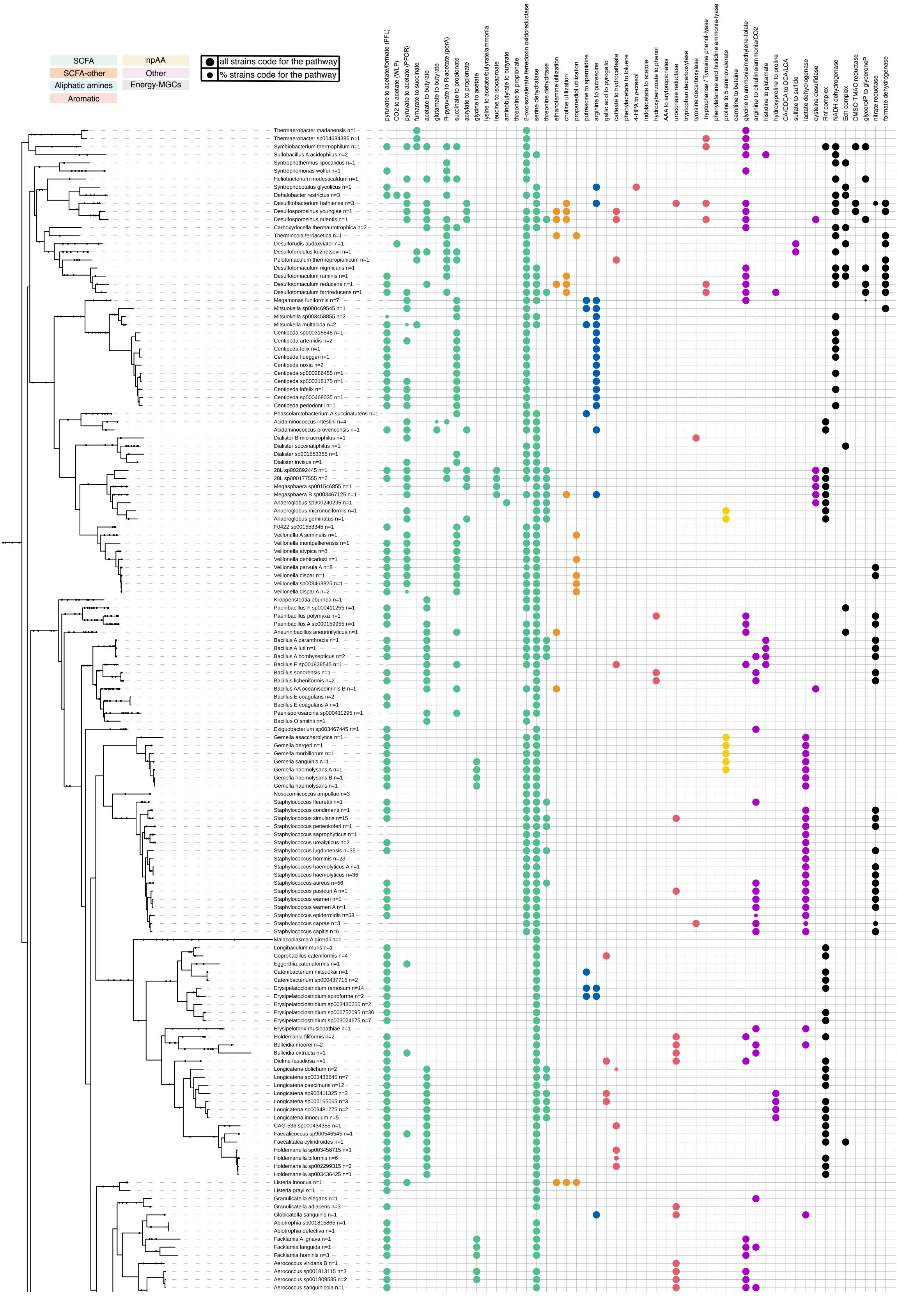

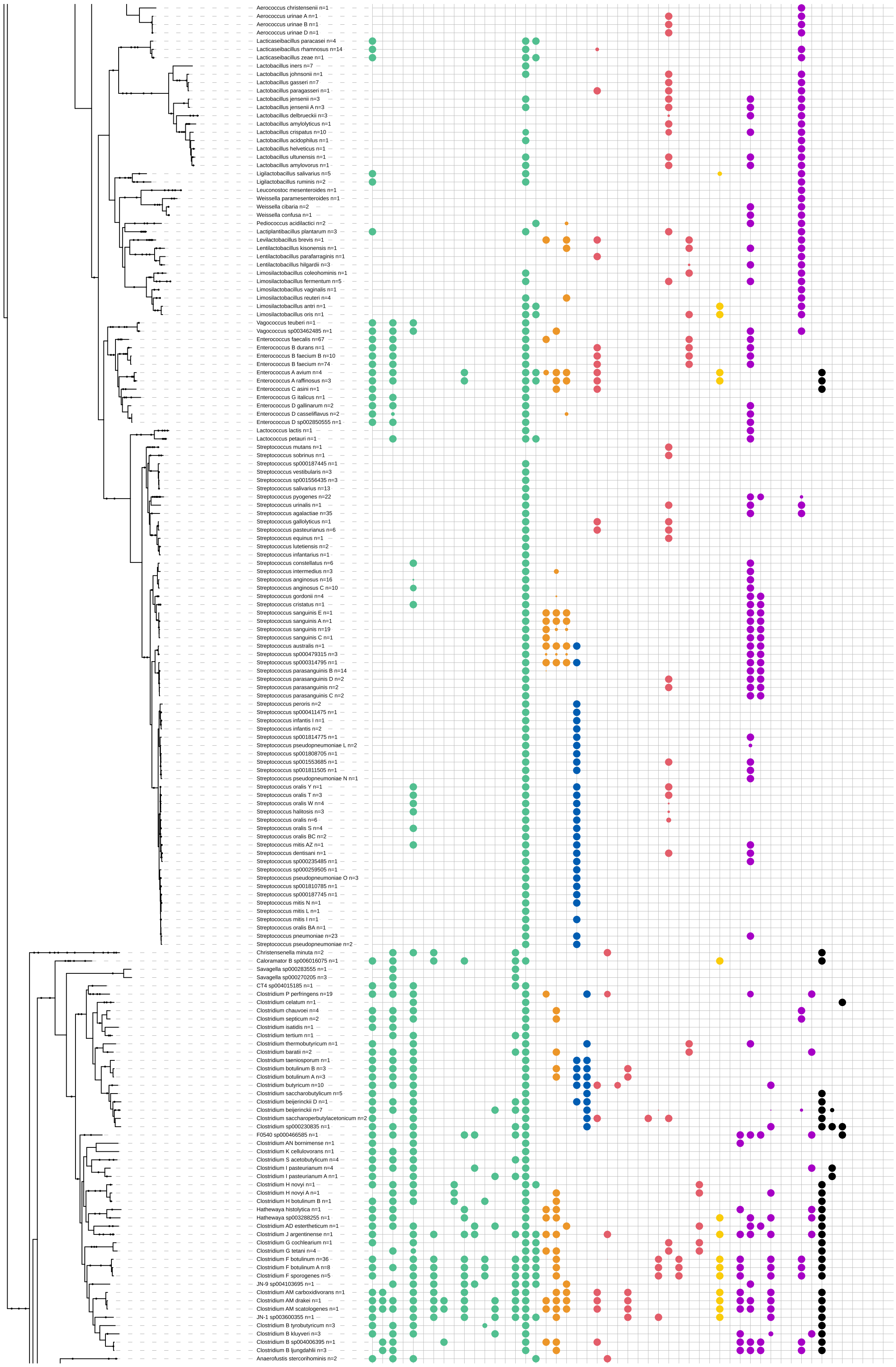

Aerococcus christensenii n=1
Aerococcus urinae A n=1
Aerococcus urinae B n=1
Aerococcus urinae D n=1
Lacticaseibacillus paracasei n=4
Lacticaseibacillus rhamnosus n=14
Lacticaseibacillus zeae n=1
Lactobacillus iners n=7
Lactobacillus johnsonii n=1
Lactobacillus gasseri n=7
Lactobacillus paragasseri n=1
Lactobacillus jensenii n=3
Lactobacillus jensenii A n=3
Lactobacillus delbrueckii n=3
Lactobacillus amylolyticus n=1
Lactobacillus crispatus n=10
Lactobacillus acidophilus n=1
Lactobacillus helveticus n=1
Lactobacillus ultunensis n=1
Lactobacillus amylovorus n=1
Ligilactobacillus salivarius n=5
Ligilactobacillus ruminis n=2
Leuconostoc mesenteroides n=1
Weissella paramesenteroides n=1
Weissella cibaria n=2
Weissella confusa n=1
Pediococcus acidilactici n=2
Lactiplantibacillus plantarum n=3
Levilactobacillus brevis n=1
Lentilactobacillus kisonensis n=1
Lentilactobacillus parafarraginis n=1
Lentilactobacillus hilgardii n=3
Limosilactobacillus coleohominis n=1
Limosilactobacillus fermentum n=5
Limosilactobacillus vaginalis n=1
Limosilactobacillus reuteri n=4
Limosilactobacillus antri n=1
Limosilactobacillus oris n=1
Vagococcus teuberi n=1
Vagococcus sp003462485 n=1
Enterococcus faecalis n=67
Enterococcus B durans n=1
Enterococcus B faecium B n=10
Enterococcus B faecium n=74
Enterococcus A avium n=4
Enterococcus A raffinosus n=3
Enterococcus C asini n=1
Enterococcus G italicus n=1
Enterococcus D gallinarum n=2
Enterococcus D casseliflavus n=2
Enterococcus D sp002850555 n=1
Lactococcus lactis n=1
Lactococcus petauri n=1
Streptococcus mutans n=1
Streptococcus sobrinus n=1
Streptococcus sp000187445 n=1
Streptococcus vestibularis n=3
Streptococcus sp001556435 n=3
Streptococcus salivarius n=13
Streptococcus pyogenes n=22
Streptococcus urinalis n=1
Streptococcus agalactiae n=35
Streptococcus gallolyticus n=1
Streptococcus pasteurianus n=6
Streptococcus equinus n=1
Streptococcus lutetiensis n=2
Streptococcus infantarius n=1
Streptococcus constellatus n=6
Streptococcus intermedius n=3
Streptococcus anginosus n=16
Streptococcus anginosus C n=10
Streptococcus gordonii n=4
Streptococcus cristatus n=1
Streptococcus sanguinis E n=1
Streptococcus sanguinis A n=1
Streptococcus sanguinis n=19
Streptococcus sanguinis C n=1
Streptococcus australis n=1
Streptococcus sp000479315 n=3
Streptococcus sp000314795 n=1
Streptococcus parasanguinis B n=14
Streptococcus parasanguinis D n=2
Streptococcus parasanguinis n=2
Streptococcus parasanguinis C n=2
Streptococcus peroris n=2
Streptococcus sp000411475 n=1
Streptococcus infantis I n=1
Streptococcus infantis n=2
Streptococcus sp001814775 n=1
Streptococcus pseudopneumoniae L n=2
Streptococcus sp001808705 n=1
Streptococcus sp001553685 n=1
Streptococcus sp001811505 n=1
Streptococcus pseudopneumoniae N n=1
Streptococcus oralis Y n=1
Streptococcus oralis T n=3
Streptococcus oralis W n=4
Streptococcus halitosis n=3
Streptococcus oralis n=6
Streptococcus oralis S n=4
Streptococcus oralis BC n=2
Streptococcus mitis AZ n=1
Streptococcus dentisani n=1
Streptococcus sp000235485 n=1
Streptococcus sp000259505 n=1
Streptococcus pseudopneumoniae O n=3
Streptococcus sp001810785 n=1
Streptococcus sp000187745 n=1
Streptococcus mitis N n=1
Streptococcus mitis L n=1
Streptococcus mitis I n=1
Streptococcus oralis BA n=1
Streptococcus pneumoniae n=23
Streptococcus pseudopneumoniae n=2
Christensenella minuta n=2
Caloramator B sp006016075 n=1
Savagella sp000283555 n=1
Savagella sp000270205 n=3
CT4 sp004015185 n=1
Clostridium P perfringens n=19
Clostridium celatum n=1
Clostridium chauvoei n=4
Clostridium septicum n=2
Clostridium isatidis n=1
Clostridium tertium n=1
Clostridium thermobutyricum n=1
Clostridium baratii n=2
Clostridium taeniosporum n=1
Clostridium botulinum B n=3
Clostridium botulinum A n=3
Clostridium butyricum n=10
Clostridium saccharobutylicum n=5
Clostridium beijerinckii D n=1
Clostridium beijerinckii n=7
Clostridium saccharoperbutylacetonicum n=2
Clostridium sp000230835 n=1
F0540 sp000466585 n=1
Clostridium AN bornimense n=1
Clostridium K cellulovorans n=1
Clostridium S acetobutylicum n=4
Clostridium I pasteurianum n=4
Clostridium I pasteurianum A n=1
Clostridium H novyi n=1
Clostridium H novyi A n=1
Clostridium H botulinum B n=1
Hathewaya histolytica n=1
Hathewaya sp003288255 n=1
Clostridium AD estertheticum n=1
Clostridium J argentinense n=1
Clostridium G cochlearium n=1
Clostridium G tetani n=4
Clostridium F botulinum n=36
Clostridium F botulinum A n=8
Clostridium F sporogenes n=5
JN-9 sp004103695 n=1
Clostridium AM carboxidivorans n=1
Clostridium AM drakei n=1
Clostridium AM scatologenes n=1
JN-1 sp003600355 n=1
Clostridium B tyrobutyricum n=3
Clostridium B kluyveri n=3
Clostridium B sp004006395 n=1
Clostridium B ljungdahlii n=3
Anaerofustis stercorihominis n=2

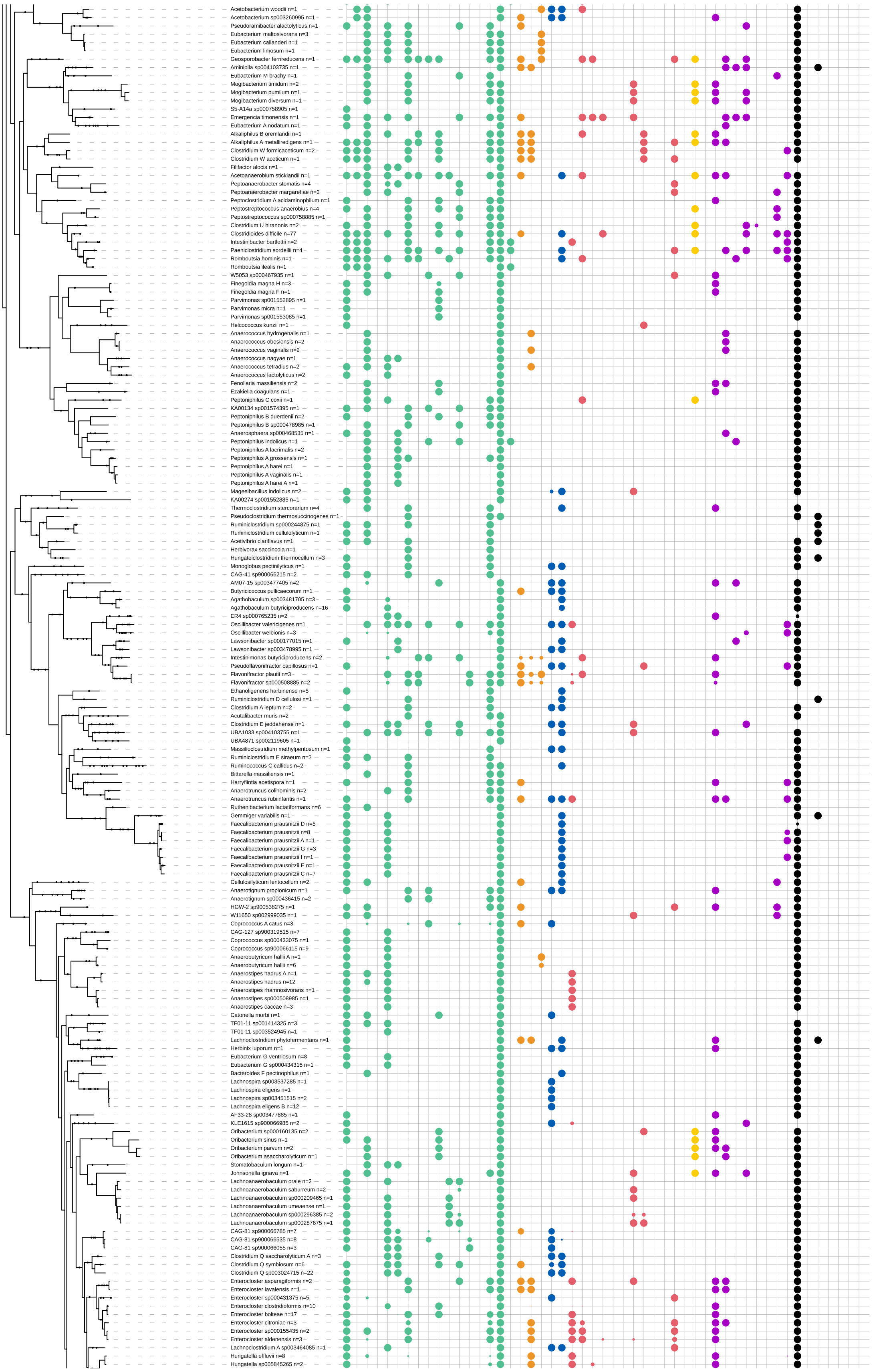
Acetobacterium woodii n=1
Acetobacterium sp003260995 n=1
Pseudoramibacter alactolyticus n=1
Eubacterium maltosivorans n=3
Eubacterium callanderi n=1
Eubacterium limosum n=1
Geosporobacter ferrireducens n=1
Aminipila sp004103735 n=1
Eubacterium M brachy n=1
Mogibacterium timidum n=2
Mogibacterium pumilum n=1
Mogibacterium diversum n=1
S5-A14a sp000758905 n=1
Emergencia timonensis n=1
Eubacterium A nodatum n=1
Alkaliphilus B oremlandii n=1
Alkaliphilus A metalliredigens n=1
Clostridium W formicaceticum n=2
Clostridium W aceticum n=1
Filifactor alocis n=1
Acetoanaerobium sticklandii n=1
Peptoanaerobacter stomatis n=4
Peptoanaerobacter margaretiae n=2
Peptoclostridium A acidaminophilum n=1
Peptostreptococcus anaerobius n=4
Peptostreptococcus sp000758885 n=1
Clostridium U hiranonis n=2
Clostridioides difficile n=77
Intestinibacter bartlettii n=2
Paeniclostridium sordellii n=4
Romboutsia hominis n=1
Romboutsia ilealis n=1
W5053 sp000467935 n=1
Finegoldia magna H n=3
Finegoldia magna F n=1
Parvimonas sp001552895 n=1
Parvimonas micra n=1
Parvimonas sp001553085 n=1
Helcococcus kunzii n=1
Anaerococcus hydrogenalis n=1
Anaerococcus obesiensis n=2
Anaerococcus vaginalis n=2
Anaerococcus nagyae n=1
Anaerococcus tetradius n=2
Anaerococcus lactolyticus n=2
Fenollaria massiliensis n=2
Ezakiella coagulans n=1
Peptoniphilus C coxii n=1
KA00134 sp001574395 n=1
Peptoniphilus B duerdenii n=2
Peptoniphilus B sp000478985 n=1
Anaerosphaera sp000468535 n=1
Peptoniphilus indolicus n=1
Peptoniphilus A lacrimalis n=2
Peptoniphilus A grossensis n=1
Peptoniphilus A harei n=1
Peptoniphilus A vaginalis n=1
Peptoniphilus A harei A n=1
Mageeibacillus indolicus n=2
KA00274 sp001552885 n=1
Thermoclostridium stercorarium n=4
Pseudoclostridium thermosuccinogenes n=1
Ruminiclostridium sp000244875 n=1
Ruminiclostridium cellulolyticum n=1
Acetivibrio clariflavus n=1
Herbivorax saccincola n=1
Hungateiclostridium thermocellum n=3
Monoglobus pectinilyticus n=1
CAG-41 sp900066215 n=1
AM07-15 sp003477405 n=2
Butyricicoccus pullicaecorum n=1
Agathobaculum sp003481705 n=3
Agathobaculum butyriciproducens n=16
ER4 sp000765235 n=2
Oscillibacter valericigenes n=1
Oscillibacter welbionis n=3
Lawsonibacter sp000177015 n=1
Lawsonibacter sp003478995 n=1
Intestinimonas butyriciproducens n=2
Pseudoflavonifractor capillosus n=1
Flavonifractor plautii n=3
Flavonifractor sp000508885 n=2
Ethanoligenens harbinense n=5
Ruminiclostridium D cellulosi n=1
Clostridium A leptum n=2
Acutalibacter muris n=2
Clostridium E jeddahense n=1
UBA1033 sp004103755 n=1
UBA4871 sp002119605 n=1
Massiliioclostridium methylpentosum n=1
Ruminiclostridium E siraeum n=3
Ruminococcus C callidus n=2
Bittarella massiliensis n=1
Harryflintia acetispora n=1
Anaerotruncus colihominis n=2
Anaerotruncus rubiinfantis n=1
Ruthenibacterium lactatiformans n=6
Gemmiger variabilis n=1
Faecalibacterium prausnitzii D n=5
Faecalibacterium prausnitzii n=8
Faecalibacterium prausnitzii A n=1
Faecalibacterium prausnitzii G n=3
Faecalibacterium prausnitzii I n=1
Faecalibacterium prausnitzii E n=1
Faecalibacterium prausnitzii C n=7
Cellulosilyticum lentocellum n=2
Anaerotignum propionicum n=1
Anaerotignum sp000436415 n=2
HGW-2 sp900538275 n=1
W11650 sp002999035 n=1
Coprococcus A catus n=3
CAG-127 sp900319515 n=7
Coprococcus sp000433075 n=1
Coprococcus sp900066115 n=9
Anaerobutyricum hallii A n=1
Anaerobutyricum hallii n=6
Anaerostipes hadrus A n=1
Anaerostipes hadrus n=12
Anaerostipes rhamnosivorans n=1
Anaerostipes sp000508985 n=1
Anaerostipes caccae n=3
Catonella morbi n=1
TF01-11 sp001414325 n=3
TF01-11 sp003524945 n=1
Lachnoclostridium phytofermentans n=1
Herbinix luporum n=1
Eubacterium G ventriosum n=8
Eubacterium G sp000434315 n=1
Bacteroides F pectinophilus n=1
Lachnospira sp003537285 n=1
Lachnospira eligens n=1
Lachnospira sp003451515 n=2
Lachnospira eligens B n=12
AF33-28 sp003477885 n=1
KLE1615 sp900066985 n=2
Oribacterium sp000160135 n=2
Oribacterium sinus n=1
Oribacterium parvum n=2
Oribacterium asaccharolyticum n=1
Stomatobaculum longum n=1
Johnsonella ignava n=1
Lachnoanaerobaculum orale n=2
Lachnoanaerobaculum saburreum n=2
Lachnoanaerobaculum sp000209465 n=1
Lachnoanaerobaculum umeaense n=1
Lachnoanaerobaculum sp000296385 n=2
Lachnoanaerobaculum sp000287675 n=1
CAG-81 sp900066785 n=7
CAG-81 sp900066535 n=8
CAG-81 sp900066055 n=3
Clostridium Q saccharolyticum A n=3
Clostridium Q symbiosum n=6
Clostridium Q sp003024715 n=22
Enterocloster asparagiformis n=2
Enterocloster lavalensis n=1
Enterocloster sp000431375 n=5
Enterocloster clostridioformis n=10
Enterocloster bolteae n=17
Enterocloster citroniae n=3
Enterocloster sp000155435 n=2
Enterocloster aldenensis n=3
Lachnoclostridium A sp003464085 n=1
Hungatella effluvii n=8
Hungatella sp005845265 n=2

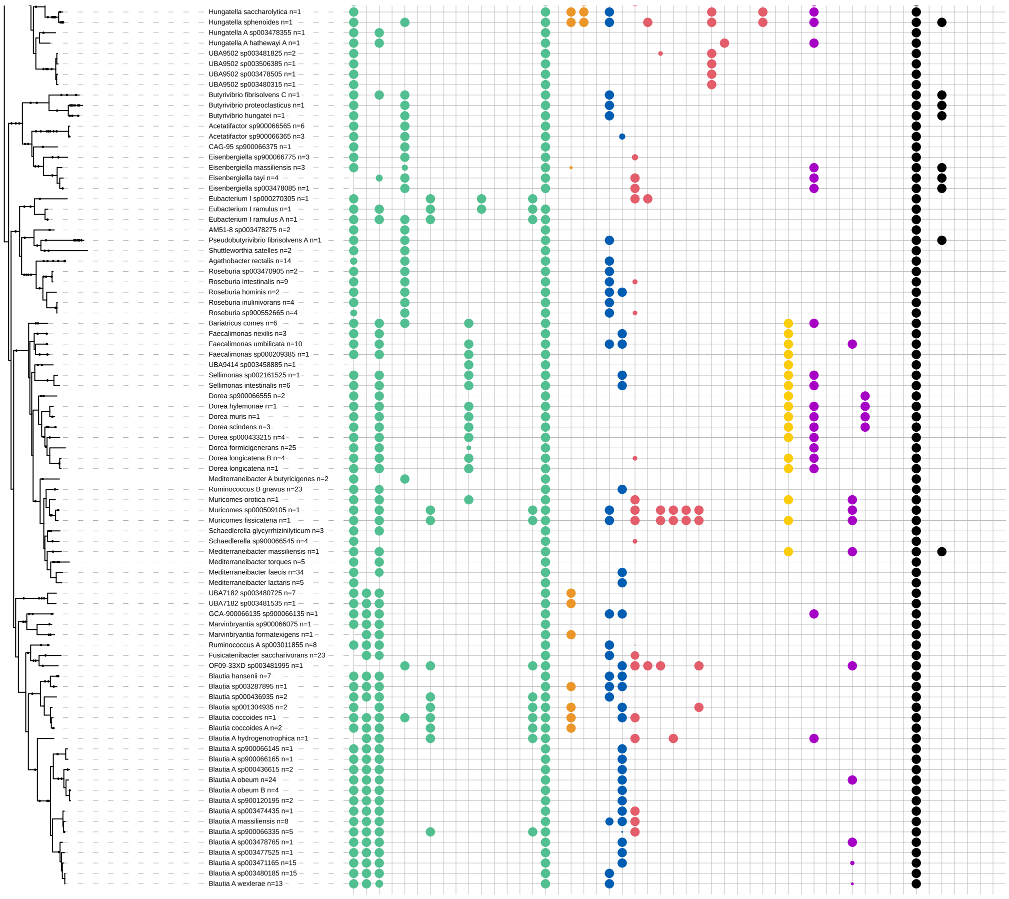


**Supplementary Figure 1: Pathway distribution across a phylogenetic tree of 557 Firmicutes present in the HMP, CGR and Clostridiales datasets.** The taxonomic assignments were performed using the GTDB database (*17*) and the phylogeny was produced using phyloT (https://phylot.biobyte.de/). Each column represents the presence/absence of the 51 metabolic pathways including single gene ones), which are color-coded based on the pathway's end product (metabolic classes). The full-sized circle implies that all strains in the node code for the pathway (see species label for information on the species group size), while smaller circles represent the relative number of species that encode the pathway. The pathways annotations were visualized using iTOL (*8*).

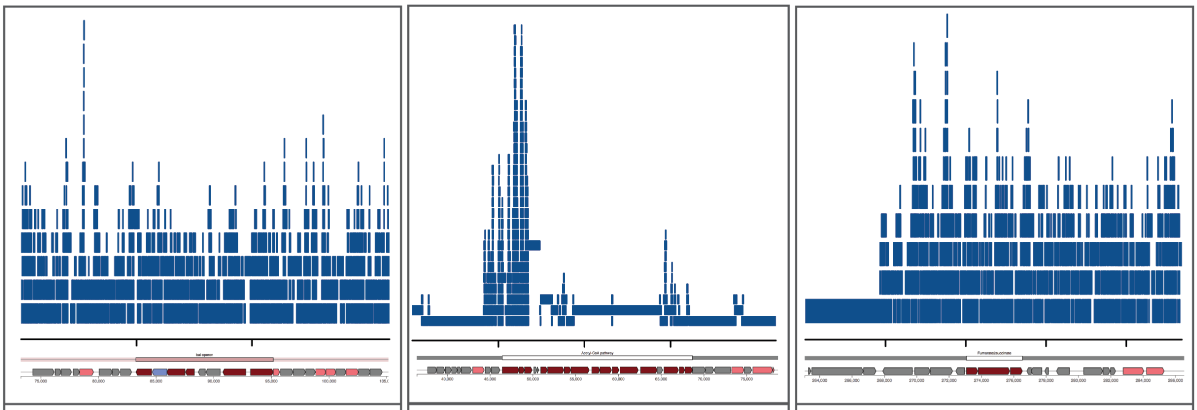

CA/CDCA to DCA/LCA *Dorea sp.* (QUDV01000004.1)
Metagenomic sample 9002000001572713

CO2 to acetate (WLP) Clostridiales bacterium (QWVV01000001)
Metagenomic sample 9002000001161852

Fumarate to succinate *Alistepes putredinis* (DS499580.1)
Metagenomic sample 9002000001302692

**Supplementary Figure 2: Read coverage of three random metagenomic samples mapping to three gene clusters of interest: the *bai* operon (CDA/CDCA to DCA/LCA), a gene cluster encoding the acetyl-CoA pathway ($CO_2$ to acetate) and a gene cluster encoding the fumarate-to-succinate pathway.** Reads are represented by blue lines, which are distributed along the x-axis based on the bedgraphs output by BiG-MAP. The plots have been produced using the Sushi R package (version 3.5.1) (*21*) and show how, despite the fact that some regions of the MGC attract more reads, the whole gene cluster is covered. They also illustrate the rationale for using the lower quartile of read coverage across 2kb regions, as this will avoid basing MGC abundance (partially) on outliers.

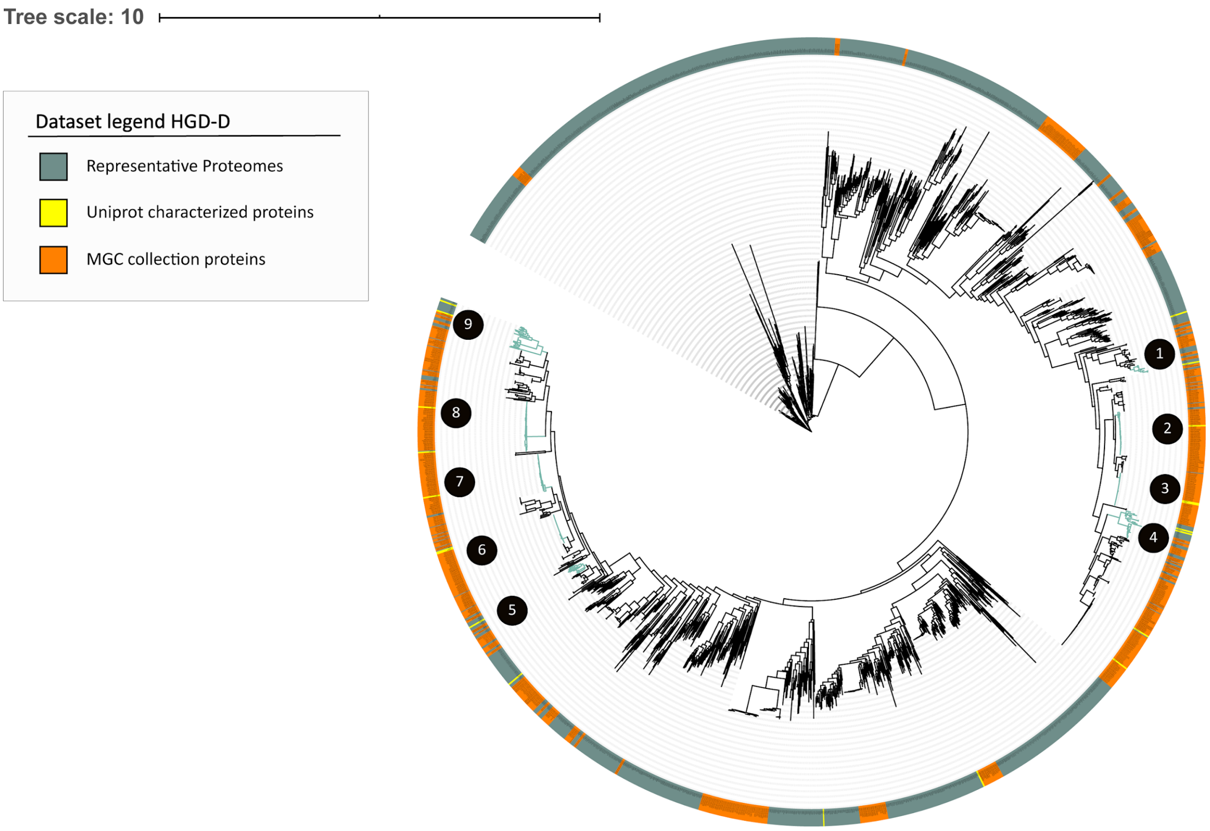


**Supplementary Figure 3: HGD-D protein superfamily phylogeny.** The phylogeny contains 2,054 protein sequences gathered from three different sources as shown with different colours in the outer ring. The highlighted clades with numbers associated are the pathway-specific clades used to create the 9 pHMMs used in the AAA to aryl propionates (AAA reductive branch), leucine to isocaproate (leucine reductive branch), glutamate to butyrate and acrylate to propionate MGC detection rules.

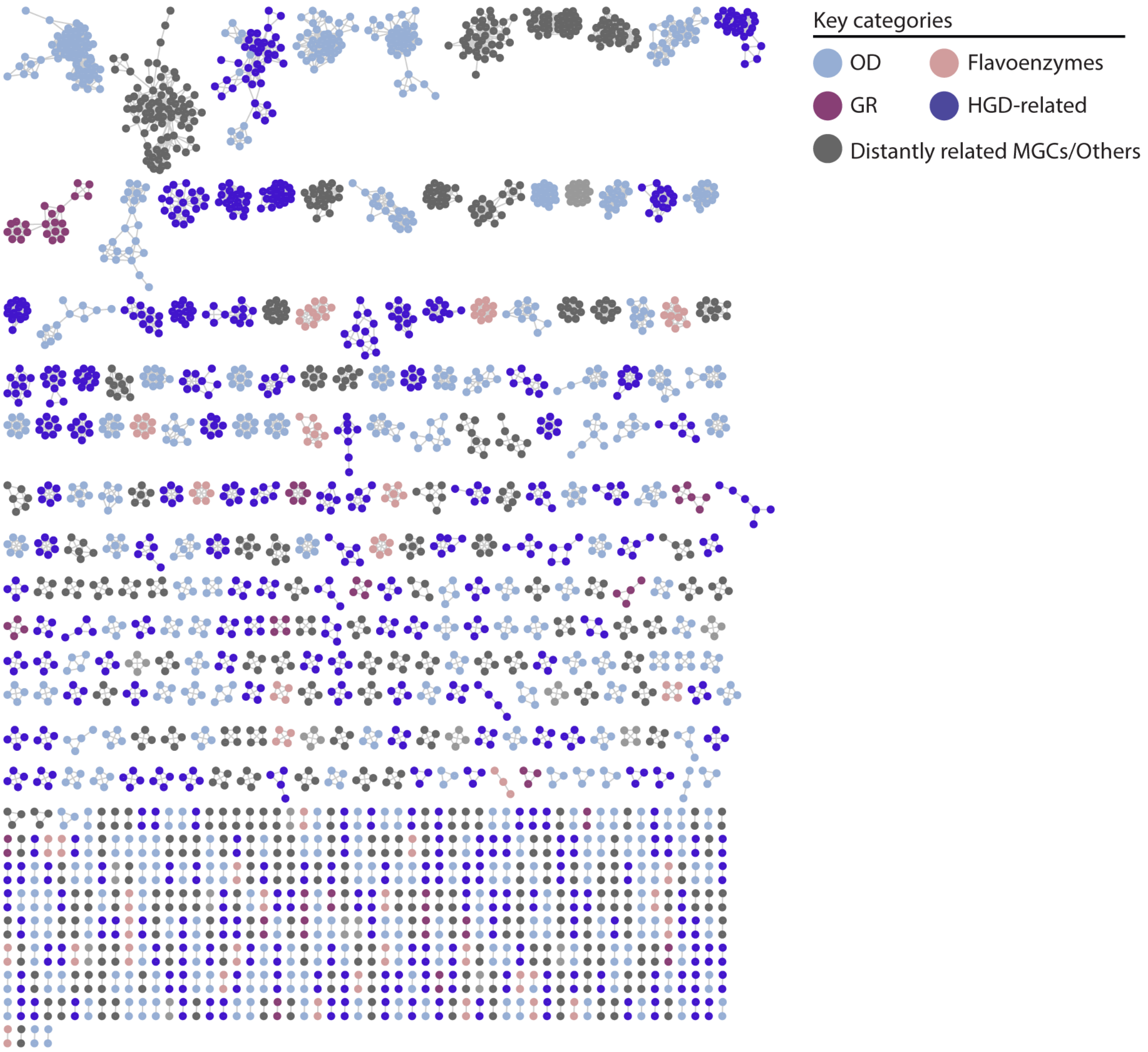


**Supplementary Figure 4: Network of putative non-redundant MGCs predicted by gutSMASH.** From all the unknown predicted MGCs, a redundancy filtering of 0.9 sequence similarity was applied using MMseqs2. From each cluster, two representatives were picked, and all representatives were used as input for BiG-SCAPE using the default cut-offs. The network contains 2,921 nodes and 7,474 edges. The MGCs have been classified into four different categories based on the key enzyme classes they code for. The GR (glycyl-radical) category is composed of MGCs that include pyruvate formate-lyase (PFL-like) and/or glycyl radical (Gly_radical), OD (oxidative decarboxylation) involves MGCs with at least one of the following Pfam domains: Pyruvate ferredoxin/flavodoxin oxidoreductase (POR), Pyruvate flavodoxin/ferredoxin oxidoreductase, thiamine diP-bdg (POR_N), Pyruvate:ferredoxin oxidoreductase core domain II (PFOR_II) and Thiamine pyrophosphate enzyme, C-terminal TPP binding domain (TPP_enzyme_C). The Flavoenzymes category is a combination of MGCs harbouring at least one of the custom-made BaiCD and BaiH pHMMs. HGD-D-related MGCs, as the name states, include enzymes matching any of the 2-hydroxyglutaryl-CoA dehydratase, D-component (HGD-D)-related pHMM domains.

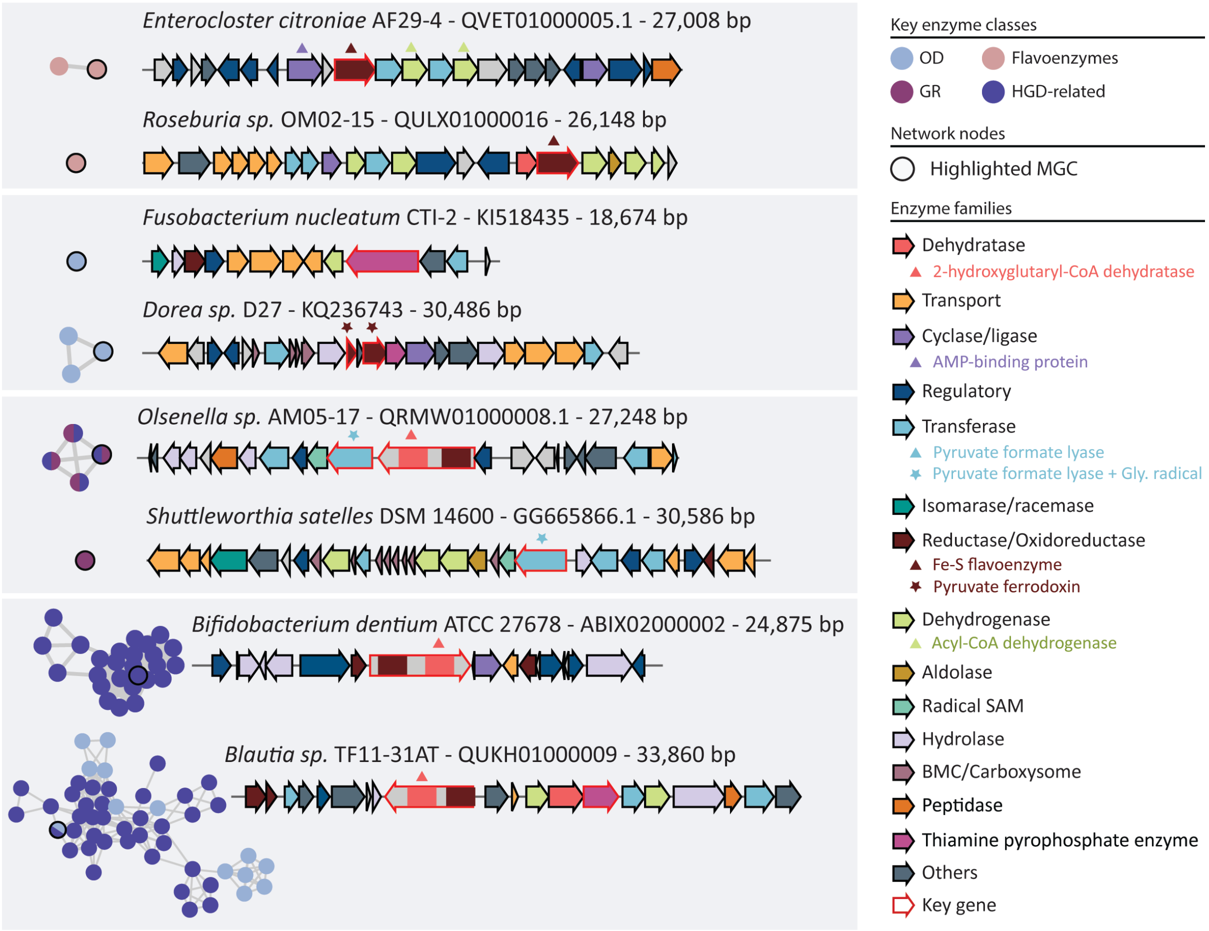


**Supplementary Figure 5: Subset of unknown MGCs predicted by gutSMASH manually picked.** The network/nodes present in the left side of the figure represent the subnetwork extracted from the complete network in Supplementary Figure 4. The arrows have been coloured-coded based on the Pfam domains found in the protein-coding sequences and the functional annotations of these proteins.

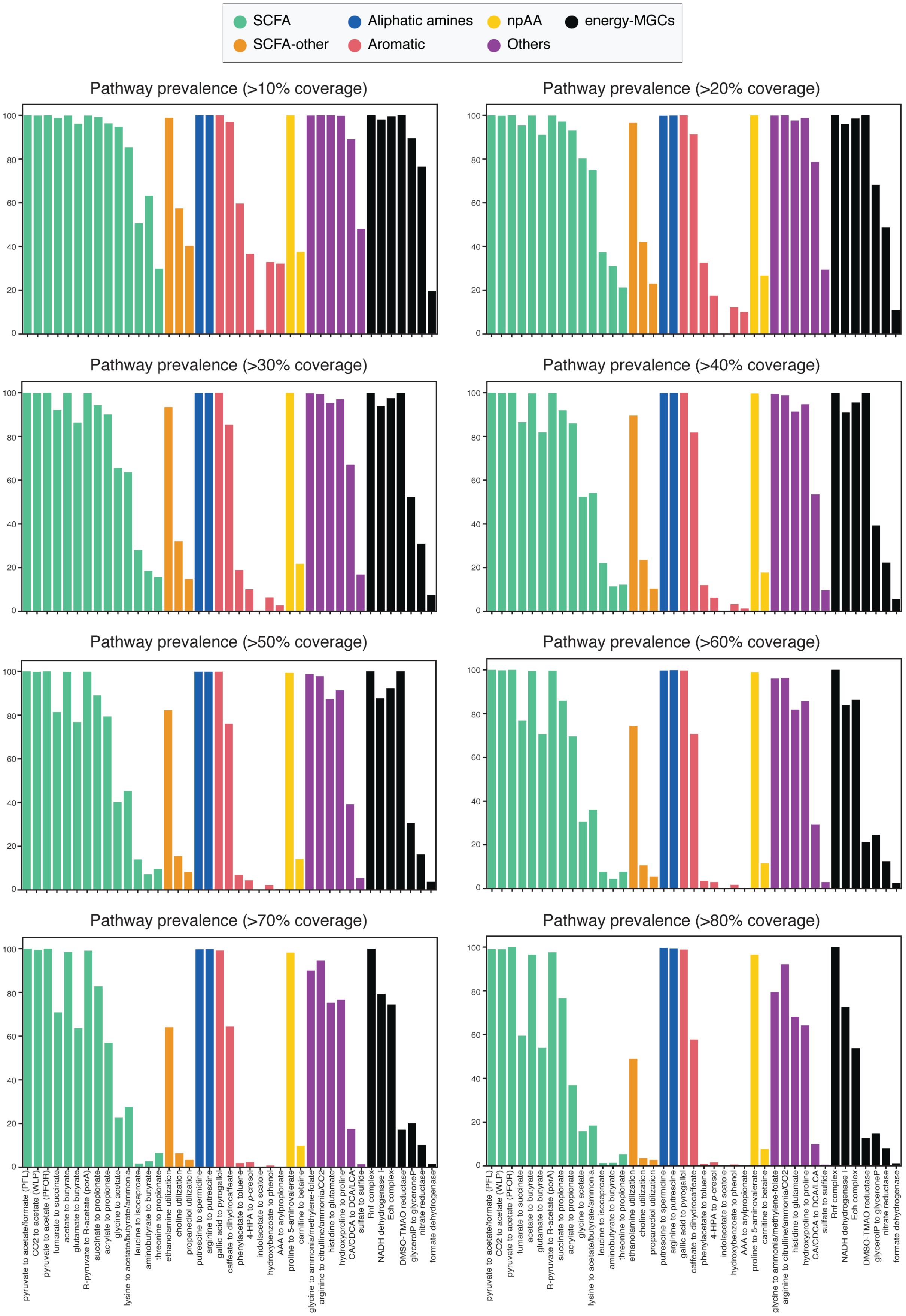
SCFA
SCFA-other
Aliphatic amines
Aromatic
npAA
Others
energy-MGCs
Pathway prevalence (>10% coverage)
Pathway prevalence (>20% coverage)
Pathway prevalence (>30% coverage)
Pathway prevalence (>40% coverage)
Pathway prevalence (>50% coverage)
Pathway prevalence (>60% coverage)
Pathway prevalence (>70% coverage)
Pathway prevalence (>80% coverage)
pyruvate to acetate/formate (PFL)
CO2 to acetate (WLP)
pyruvate to acetate (PFOR)
fumarate to succinate
acetate to butyrate
glutamate to butyrate
R-pyruvate to R-acetate (porA)
succinate to propionate
acrylate to propionate
glycine to acetate
lysine to acetate/butyrate/ammonia
leucine to isocaproate
aminobutyrate to butyrate
threonine to propionate
ethanolamine utilization
choline utilization
propanediol utilization
putrescine to spermidine
arginine to putrescine
gallic acid to pyrogallol
caffeate to dihydrocaffeate
phenylacetate to toluene
4-HPA to p-cresol
indoleacetate to scatole
hydroxybenzoate to phenol
AAA to arylpropionate
proline to 5-aminovalerate
carnitine to betaine
glycine to ammonia/methylene-folate
arginine to citrulline/ammonia/CO2
histidine to glutamate
hydroxyproline to proline
CA/CDCA to DCA/LCA
sulfate to sulfide
Rnf complex
NADH dehydrogenase I
Ech complex
DMSO-TMAO reductase
glycerolP to glyceroneP
nitrate reductase
formate dehydrogenase

**Supplementary Figure 6: Pathway prevalence using different core coverage thresholds.** Pathway prevalence was computed by assessing the number of reads (per sample) mapping to known gene clusters at a certain core coverage cut-off. The figure illustrates how the pathway prevalence gradually changes when increasing the core coverage cut-off from 10 to 80%.

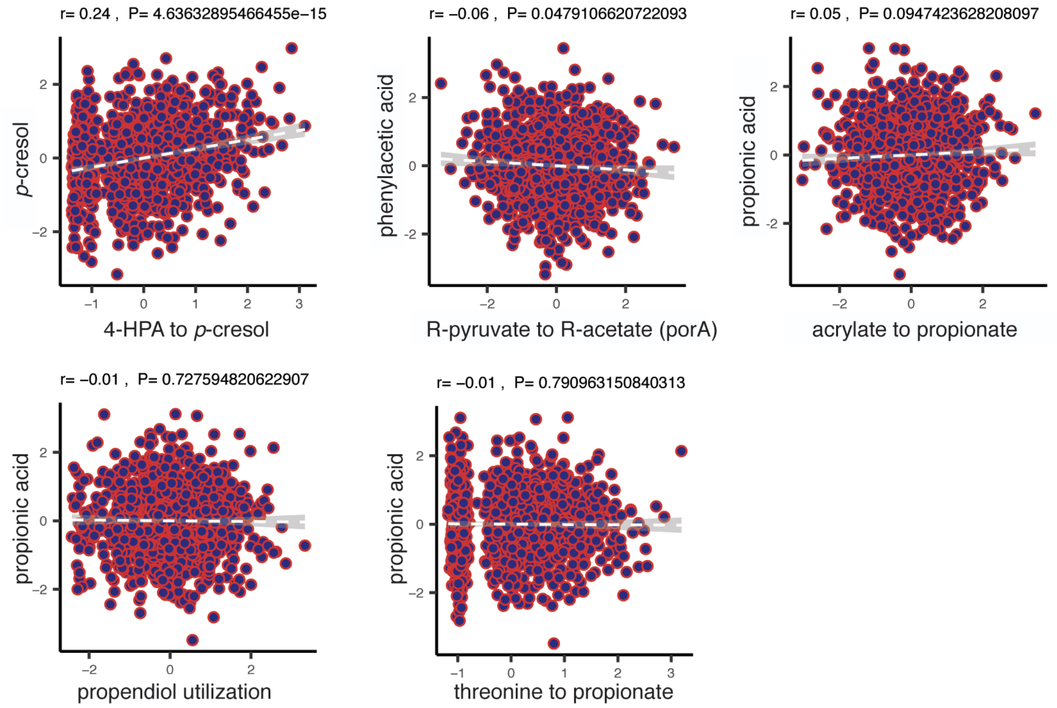


**Supplementary Figure 7: Limited correlation of genetic pathway abundance with metabolites abundance in blood plasma.** This figure shows correlation plots for additional metabolites not shown in Figure 3c.

#### Supplementary Table Legends

**Table S1: Dataset of 51 known pathways, including single-protein pathways.** This dataset helped designing detection rules for gutSMASH and build pHMMs for more accurate pathway identification. The table includes information on the substrate(s) and product(s) of each pathway, Pfams involved (ID and description) and the original organism where it has been characterized.

**Table S2: Validation results of the gutSMASH predictive potential using PaperBLAST.** Organisms whose genomes encode homologues of proteins from the original data set of known pathways were found using paperBLAST and their genomes were used as input for gutSMASH. The table includes information on the original organisms, validation organism, amino acid sequence identity and overall sequence identity.

**Table S3: Assembly IDs of the 1,621 genomes used to validate the gutSMASH detection rules.**

**Table S4: Assembly IDs of 4,240 genomes from the HMP, CGR and Clostridiales collections used to screen the metabolic potential of human gut bacteria.**

**Table S5: Raw pathway abundance across most representative genera of bacteria in the human gut.** This data has been used to create Figure 2a.

**Table S6: Absolute counts of genomes harboring genes and MGCs corresponding to the main acetate-producing pathways, summarized at phylum level.** This data has been used to create Figure 2b.

**Table S7: MetaCyc links found for 38/41 of the pathways predicted by gutSMASH.**

**Table S8: Pathway prevalence values of the 41 pathways across 1,135 human microbiomes using different BiG-MAP mapping coverage threshold values.** These values have been used to create figure 3a and Supplementary Figure 6.

**Table S9: Pathway abundance counts across 1,135 human microbiome samples.** These values have been used to create Figure 3b.

**Table S10: List of pHMMs created to accurately identify pathway-specific clades of interest.** These pHMMs have been included in the gutSMASH v1.0 detection rule set and are available online at https://github.com/victoriapascal/gutsmash/tree/gutsmash/antismash/detection/gut_hmm_detection/data .

**Table S11: Overview of detection rules in gutSMASH v1.0 for the identification of MGCs encoding known pathways.**

**Table S12: Set of known pathways used for the iterative homology search approach.** The ultimate goal of this approach was to design the general rules. The original Genbank files can be downloaded from https://gutsmash.bioinformatics.nl/help.html#Pathways-subset .

**Table S13: Overview of general detection rules used by gutSMASH version 1.0.**

#### References


1. C. Chen, D. A. Natale, R. D. Finn, H. Huang, J. Zhang, C. H. Wu, R. Mazumder, Representative proteomes: a stable, scalable and unbiased proteome set for sequence analysis and functional annotation. *PLoS ONE* **6**, e18910 (2011).

2. M. N. Price, A. P. Arkin, PaperBLAST: text mining papers for information about homologs. *mSystems*. **2** , mSystems.00039-17 (2017).

3. J. C. Navarro-Muñoz, N. Selem-Mojica, M. W. Mullowney, S. A. Kautsar, J. H. Tryon, E. I. Parkinson, E. L. C. De Los Santos, M. Yeong, P. Cruz-Morales, S. Abubucker, A. Roeters, W. Lokhorst, A. Fernandez-Guerra, L. T. D. Cappelini, A. W. Goering, R. J. Thomson, W. W. Metcalf, N. L. Kelleher, F. Barona-Gomez, M. H. Medema, A computational framework to explore large-scale biosynthetic diversity. *Nat. Chem. Biol.* **16**, 60–68 (2020).

4. E. F. Tigchelaar, A. Zhernakova, J. A. M. Dekens, G. Hermes, A. Baranska, Z. Mujagic, M. A. Swertz, A. M. Muñoz, P. Deelen, M. C. Cénit, L. Franke, S. Scholtens, R. P. Stolk, C. Wijmenga, E. J. M. Feskens, Cohort profile: LifeLines DEEP, a prospective, general population cohort study in the northern Netherlands: study design and baseline characteristics. *BMJ Open* **5** (2015), e006772.

5. A. Zhernakova, A. Kurilshikov, M. J. Bonder, E. F. Tigchelaar, M. Schirmer, T. Vatanen, Z. Mujagic, A. V. Vila, G. Falony, S. Vieira-Silva, J. Wang, F. Imhann, E. Brandsma, S. A. Jankipersadsing, M. Joossens, M. C. Cenit, P. Deelen, M. A. Swertz, LifeLines cohort study, R. K. Weersma, E. J. M. Feskens, M. G. Netea, D. Gevers, D. Jonkers, L. Franke, Y. S. Aulchenko, C. Huttenhower, J. Raes, M. H. Hofker, R. J. Xavier, C. Wijmenga, J. Fu, Population-based metagenomics analysis reveals markers for gut microbiome composition and diversity. *Science* **352**, 565–569 (2016).

6. F. Sievers, A. Wilm, D. Dineen, T. J. Gibson, K. Karplus, W. Li, R. Lopez, H. McWilliam, M. Remmert, J. Söding, J. D. Thompson, D. G. Higgins, Fast, scalable generation of high-quality protein multiple sequence alignments using Clustal Omega. *Mol. Syst. Biol.* **7**, 539 (2011).

7. M. N. Price, P. S. Dehal, A. P. Arkin, FastTree 2 – approximately maximum-likelihood trees for large alignments. *PLoS ONE* **5**, e9490 (2010).

8. I. Letunic, P. Bork, Interactive Tree Of Life (iTOL) v4: recent updates and new developments. *Nucleic Acids Res.* **47**, W256–W259 (2019).

9. A. M. Waterhouse, J. B. Procter, D. M. A. Martin, M. Clamp, G. J. Barton, Jalview version 2--a multiple sequence alignment editor and analysis workbench. *Bioinformatics*. **25**, 1189–1191 (2009).

10. Y. Zou, W. Xue, G. Luo, Z. Deng, P. Qin, R. Guo, H. Sun, Y. Xia, S. Liang, Y. Dai, D. Wan, R. Jiang, L. Su, Q. Feng, Z. Jie, T. Guo, Z. Xia, C. Liu, J. Yu, Y. Lin, S. Tang, G. Huo, X. Xu, Y. Hou, X. Liu, J. Wang, H. Yang, K. Kristiansen, J. Li, H. Jia, L. Xiao, 1,520 reference genomes from cultivated human gut bacteria enable functional microbiome analyses. *Nat. Biotechnol.* **37**, 179–185 (2019).

11. C. Camacho, G. Coulouris, V. Avagyan, N. Ma, J. Papadopoulos, K. Bealer, T. L. Madden, BLAST+: architecture and applications. *BMC Bioinformatics*. **10**, 421 (2009).

12. V. Pascal Andreu, M. A. Fischbach, M. H. Medema, Computational genomic discovery of diverse

gene clusters harbouring Fe-S flavoenzymes in anaerobic gut microbiota. *Microb. Genom*. **6**, mgen.0.000373 (2020).

13. E. L. C. de los Santos, G. L. Challis, clusterTools: proximity searches for functional elements to identify putative biosynthetic gene clusters. *Access Microbiol.* **1**, po0154 (2019).

14. M. H. Medema, E. Takano, R. Breitling, Detecting sequence homology at the gene cluster level with MultiGeneBlast. *Mol. Biol. Evol.* **30**, 1218–1223 (2013).

15. M. Steinegger, J. Söding, MMseqs2 enables sensitive protein sequence searching for the analysis of massive data sets. *Nat. Biotechnol.* **35**, 1026–1028 (2017).

16. D. Hyatt, G.-L. Chen, P. F. Locascio, M. L. Land, F. W. Larimer, L. J. Hauser, Prodigal: prokaryotic gene recognition and translation initiation site identification. *BMC Bioinformatics*. **11**, 119 (2010).

17. D. H. Parks, M. Chuvochina, D. W. Waite, C. Rinke, A. Skarshewski, P.-A. Chaumeil, P. Hugenholtz, A standardized bacterial taxonomy based on genome phylogeny substantially revises the tree of life. *Nat. Biotechnol.* **36**, 996–1004 (2018).

18. P. Shannon, A. Markiel, O. Ozier, N. S. Baliga, J. T. Wang, D. Ramage, N. Amin, B. Schwikowski, T. Ideker, Cytoscape: a software environment for integrated models of biomolecular interaction networks. *Genome Res.* **13**, 2498–2504 (2003).

19. V. P. Andreu, H. E. Augustijn, K. van den Berg, J. J. J. van der Hooft, M. A. Fischbach, M. H. Medema, BiG-MAP: an automated pipeline to profile metabolic gene cluster abundance and expression in microbiomes. *BioRxiv*, doi:10.1101/2020.12.14.422671 (2020).

20. B. D. Ondov, T. J. Treangen, P. Melsted, A. B. Mallonee, N. H. Bergman, S. Koren, A. M. Phillippy, Mash: fast genome and metagenome distance estimation using MinHash. *Genome Biol.* **17**, 132 (2016).

21. D. H. Phanstiel, A. P. Boyle, C. L. Araya, M. P. Snyder, Sushi.R: flexible, quantitative and integrative genomic visualizations for publication-quality multi-panel figures. *Bioinformatics*. **30**, 2808–2810 (2014).
